# Deletion of calcineurin from GFAP-expressing astrocytes impairs excitability of cerebellar and hippocampal neurons through astroglial Na^+^/K^+^ ATPase

**DOI:** 10.1101/728741

**Authors:** Laura Tapella, Teresa Soda, Lisa Mapelli, Valeria Bortolotto, Heather Bondi, Federico A Ruffinatti, Giulia Dematteis, Alessio Stevano, Marianna Dionisi, Simone Ummarino, Annalisa Di Ruscio, Carla Distasi, Mariagrazia Grilli, Armando A Genazzani, Egidio D’Angelo, Francesco Moccia, Dmitry Lim

**Author notes:** These Authors contributed equally (Laura Tapella, Teresa Soda). Correspondence should be sent to: Egidio D’Angelo; Dmitry Lim. **Competing interests** The authors have no competing interests. The data that support the findings of this study are available from the corresponding author upon reasonable request.

## Abstract

Astrocytes perform important housekeeping functions in the nervous system including maintenance of adequate neuronal excitability, although the regulatory mechanisms are currently poorly understood. The astrocytic Ca^2+^/calmodulin-activated phosphatase calcineurin (CaN) is implicated in the development of reactive gliosis and neuroinflammation, but its roles, including the control of neuronal excitability, in healthy brain is unknown. We have generated a mouse line with conditional knockout (KO) of CaN B1 (CaNB1) in glial fibrillary acidic protein (GFAP)-expressing astrocytes (<u>a</u>stroglial <u>c</u>alci<u>n</u>eurin <u>k</u>nock-<u>o</u>ut, ACN-KO). Here we report that postnatal and astrocyte-specific ablation of CaNB1 did not alter normal growth and development as well as adult neurogenesis. Yet, we found that specific deletion of astrocytic CaN selectively impairs intrinsic neuronal excitability in hippocampal CA1 pyramidal neurons and cerebellar granule cells (CGCs). This impairment was associated with a decrease in after-hyperpolarization in CGC, while passive properties were unchanged, suggesting impairment of K^+^ homeostasis. Indeed, blockade of Na^+^/K^+^-ATPase (NKA) with ouabain phenocopied the electrophysiological alterations observed in ACN-KO CGCs. In addition, NKA activity was significantly lower in cerebellar and hippocampal lysates and in pure astrocytic cultures from ACN-KO mice. While no changes were found in protein levels, NKA activity was inhibited by the specific CaN inhibitor FK506 in both cerebellar lysates and primary astroglia from control mice, suggesting that CaN directly modulates NKA activity and in this manner controls neuronal excitability. In summary, our data provide formal evidence for the notion that astroglia is fundamental for controlling basic neuronal functions and place CaN center-stage as an astrocytic Ca^2+^-sensitive switch.

## 1. INTRODUCTION

The Ca^2+^/calmodulin (CaM)-activated serine-threonine phosphatase, calcineurin (CaN), is abundantly expressed in neurons (Klee et al., 1979) but also in other brain cellular types, including astrocytes (Lim et al., 2016). CaN is a heterodimer composed of a larger (59 kDa) catalytic subunit A (CaNA) and a smaller (19 kDa) obligatory regulatory subunit B (CaNB). Three genes (Ppp3ca, Ppp3cb and Ppp3cc) encode for CaNA (α, β and γ) and two genes (Ppp3r1 and Ppp3r2) encode for CaNB (1 and 2). Two catalytic (CaNAα and CaNAβ) and one regulatory (CaNB1) subunits are expressed in the brain and therefore the selective suppression of CaN activity in mouse models is routinely achieved by deletion of the regulatory CaNB1 subunit (Zeng et al., 2001; Miyakawa et al., 2003).

CaN is activated by an increase in cytosolic Ca^2+^ concentrations ([Ca^2+^]_cyt_) and it has been shown that low but long-lasting elevations of [Ca^2+^]_cyt_ specifically activate CaN (Negulescu et al., 1994; Dolmetsch et al., 1997). Neuronal CaN plays a crucial role in the control of neurotransmission, synaptic plasticity and memory formation (Zeng et al., 2001; Baumgärtel et al., 2008). Accordingly, it has long been known that CaN finely regulates the strength of synaptic transmission by eliciting long-term depression (LTD) and by constraining long-term potentiation (LTP) or resetting potentiated synapses (Baumgärtel and Mansuy, 2012). In addition, endogenous CaN may modulate neuronal excitability. For instance, an increase in CaN activity reduces the firing rate by up-regulating the A-type K^+^ channel, Kv4.2 in mouse pyramidal neurons (Yao et al., 2016) and by shortening the axon initial segment in mouse dentate granule cells (Evans et al., 2015). Although expressed at significantly lower levels in astrocytes compared to neurons, astrocytic CaN is upregulated and plays a crucial role in astrogliosis and neuroinflammation (Pekny et al., 2016). For instance, astroglial CaN regulates neuronal excitability in a murine model of Alzheimer’s Disease (Sompol et al., 2017) and modulates synaptic strength and long-term plasticity in a rat model of traumatic brain injury (Furman et al., 2016). However, there is no evidence about the role of astroglial CaN in neuronal excitability in the healthy brain.

Here, we report the generation and characterization of a mouse model with conditional CaNB1 KO specific for glial fibrillary acidic protein (GFAP)-expressing astrocytes (<u>a</u>stroglial <u>c</u>alci<u>n</u>eurin <u>k</u>nock-<u>o</u>ut, ACN-KO). We demonstrate that ACN-KO mice do not present alterations of development, growth, brain weight and architecture and therefore are suitable for studying the role of astrocytic CaN in healthy brain. We further report that ACN-KO mice display an impairment of excitability of cerebellar granule cells (CGCs) and CA1 hippocampal pyramidal neurons and provide evidence that this may be due to the functional inactivation of the astroglial Na^+^/K^+^ ATPase. These findings provide the first evidence that astrocytic CaN modulates neuronal excitability in the healthy brain, a novel mechanism which bears remarkable consequences on information processing at the tripartite synapse.

## 2. MATERIALS AND METHODS

### 2.1. Generation and handling of ACN-KO mice

Conditional CaN knockout (KO) in astroglial cells was generated by crossing CaNB1^flox/flox^ mice (Neilson et al., 2004) (Jackson Laboratory strain B6;129S-Ppp3r1tm2Grc/J, stock number 017692), in which 3^rd^, 4^th^ and 5^th^ exons in the CaNB1, a regulatory CaN subunit gene, were flanked by LoxP sequences, with mice in which expression of bacterial Cre recombinase was controlled by the full-length murine GFAP promoter (Gregorian et al., 2009) (Jackson Laboratory strain B6.Cg-Tg(Gfap-cre)77.6Mvs/2J, stock number 024098). F1 founder mice were backcrossed for four generations and the line was maintained by crossing CaNB1^flox/flox^/GfapCre^−/−^ (thereafter referred as to ACN-Ctr) males with CaNB1^flox/flox^/GfapCre^+/−^ (ACN-KO) females (Suppl. Fig. 1A).

To generated ACN-KO mice expressing tdTomato reporter, ACN-KO mice were crossed with Rosa26-CAG-flox-stop-flox-tdTomato (RATO) mice (kind donation from Prof. Frank Kirchhoff, CIPMM, Center for Integrative Physiology and Molecular Medicine, Molecular Physiology, University of Saarland, Homburg) resulting RATO^flox/flox^/CaNB1^flox/flox^/GfapCre^+/−^ mice (thereafter referred as to RATO-ACN-KO) and RATO^flox/flox^/CaNB1^wt/wt^/GfapCre^+/−^ (RATO-ACN-Ctr) (Suppl. Fig. 1B). Genotyping was performed by means of PCR and real-time PCR. Genotyping protocols are provided in Supplementary Methods section.

Mice were housed in the animal facility of the Università del Piemonte Orientale, were kept at three to four per cage, and had unlimited access to water and food. Animals were managed in accordance with European directive 2010/63/UE and with Italian law D.l. 26/2014. The procedures were approved by the local animal-health and ethical committee (Università del Piemonte Orientale) and were authorized by the national authority (Istituto Superiore di Sanità; authorization numbers N. 77-2017 and N. 214-2019). All efforts were made to reduce the number of animals by following the 3R’s rule.

### 2.2. Histochemistry

Ctr and ACN-KO mice were perfused transcardially, brains were collected, and processed as previously described (Denis-Donini et al., 2008). Sagittal 40-µm-thick cryosections have been prepared and used for immunostaining. The following primary antibodies were used: goat anti-doublecortin (DCX) antibody (1:300; Santa Cruz Biotechnology, Santa Cruz, CA), rat anti-BrdU antibody (1:200; Novus Biologicals Inc., Littleton, CO), mouse anti-neuronal nuclei (NeuN) antibody (1:150; Millipore Corp., Billerica, MA).

#### 2.2.1. Neurogenesis assessment

DCX immunolabelling has been performed as previously described (Dellarole and Grilli, 2008) and 3 sections/mouse (n = 4/group) have been selected, spaced 640 µm in order to analyze different levels along the entire extension of dentate gyrus formation.

To analyze the neuronal differentiation of new-born cells, ACN-Ctr and ACN-KO mice (n=5/group) were intraperitoneally injected with bromodeoxyuridine (BrdU) (150 mg/kg of body weight i.p.; Sigma-Aldrich, Saint Louis, MO) for three consecutive days. Mice were sacrificed thirty days after the last BrdU injection. For immunofluorescence with antibodies against BrdU and NeuN a series of one-in-eight sections through the dentate gyrus was selected and analyzed as previously described (Stagni et al., 2017).

#### 2.2.2 Assessment of tdTomato reporter expression in neural progenitor cells and mature neurons

One brain section of ACN-Ctr and ACN-KO mice at P7 or at 1 mo of age was selected for the analysis of tdTomato reporter expression in neural progenitor cells and mature neurons, respectively. The following primary antibodies were used: rabbit anti-SOX2 antibody (1:800; Millipore), rabbit anti-NeuN antibody (1:200, Thermo Fisher). Immunolabeling was performed as previously described (Meneghini et al., 2010); then, sections were scanned with a Zeiss 710 confocal laser scanning microscope. Sox2 and NeuN positive cells were analyzed in their entire z-axis to assess the co-expression with the tdTomato reporter.

#### 2.2.3. Anti-CaNB1 staining

For anti-CaNB1 immunohistochemistry, cryosections were first subject to antigen retrieval in citrate buffer (10 mM Citric Acid, 0.05% Tween 20, pH 6.0) for 20 min at 95-100%, and probed with rabbit anti-CaNB1 antibody (1:200; Millipore Corp. Cat. 07-1494). Chicken anti-rabbit Alexa Fluor 488 secondary antibody was used. Confocal images were taken using Zeiss 710 confocal laser scanning microscope equipped with N-Achroplan 10x/0.25 or EC Plan-Neofluar 40x/1.30 Oil DIC M27 objectives and Zen software. Standard deviation of the intensity profile of anti-CaNB1 stained sections was calculated using MetaMorph software (Molecular Devices, Sunnyvale, CA, US).

### 2.3. Dissociation and FACS isolation of astrocytes

For isolation of astrocytes by FACS, P20 ACN-KO and ACN-Ctr pups were anesthetized and sacrificed by decapitation. Cerebella were dissected in cold HBSS (Sigma, Cat. H6648), minced and digested in 0.125% Trypsin (Sigma, Cat. T3924) in presence of 20 u of DNase (Sigma, Cat, D5025) at 37°C with gentle agitation for 30 min. Cerebella from 2 animals were processed in a single 2 ml Eppendorf tube. After brief centrifugation (3 min 300 xg) tissues were resuspended in 1.5 ml HBSS supplemented with 3% fetal bovine serum (FBS, Gibco, Cat. 10270) and 5 mM EDTA (HBSS-FBS-EDTA), and gently triturated with 15-20 strokes of fire polished Pasteur pipettes of three decreasing sizes. After each round of trituration the suspension was left for 2 min and 1 ml was transferred into a new 15 ml Falcon tube and fresh HBSS-FBS-EDTA was added. After three rounds of trituration, the suspension was filtered through 40 μm nylon strainer (Corning, Cat. 352340). The cells were centrifuged twice at 133 xg for 8 min and resuspended in 500 μL HBSS-FBS-EDTA. For immunostaining, 100 μL aliquots were stained with Draq5 (2 μM, Thermo Fisher, Milan, Italy, Cat. 62251), anti GLAST(ACSA-1) PE-conjugated antibody (1:250, Cat 130-118-483) and with anti-CD90.2 FITC-conjugated antibody (Cat. 130-102-452) (last two from Miltenyi Biotech, Bologna, Italy). Staining was performed at 4°C for 20 min in dark. After staining cells were washed by 1 ml of HBSS-FBS-EDTA, passed through 40 μM strainer and analyzed on S3e Cell Sorter (Bio-Rad, Segrate, Italy). Gating parameters were: singlets → Draq5^+^ (FL4)→ GLAST-PE^+^ or tdTomato^+^ (FL2) → CD90.2^−^ (FL1) (Suppl. Figure 3). From 4×10^4^ to 2×10^5^ cells were routinely sorted from one preparation directly in Trizol-LS (Invitrogen, Cat. 10296028).

To assess CnB1 expression in RATO-ACN-KO astrocytes, hippocampal tissue was dissociated enzymatically and mechanically as described above and, after enrichment by three rounds of centrifugation at 133 xg for 8 min, the cell suspension was plated on Poly-L-lysine-coated (Cat. P2636, Sigma) 13 mm round coverslips. The cell suspension was bathed in a complete culture medium (Dulbecco’s modified Eagle’s medium (DMEM, Cat. D5671), supplemented with 10% fetal bovine serum (FBS), and with 2 mg/mL glutamine, 10 U/mL penicillin and 100 mg/mL streptomycin (all from Sigma)). After 48 h of culture, cells were fixed in 4% formaldehyde in PBS, permeabilized (0.1% of Triton X-100 in PBS for 7 min at RT) and stained with primary rabbit anti-CaNB1 antibody (1:200; Millipore Corp. Cat. 07-1494) followed by chicken anti-rabbit Alexa Fluor 488 secondary antibody.

### 2.4. Primary astroglial and mixed cultures

Primary cultures were prepared as described previously (Tapella et al., 2018) with slight modifications. In brief, for purified astroglial cultures, cerebella were dissected from 3-to 5-day-old pups in cold HBSS, digested in 0.25% trypsin (37 °C, 20 min), washed in complete culture medium and resuspended in cold HBSS supplemented with 10% FBS. After 30 strokes of dissociation with an automatic pipette, cell suspension was centrifuged (250 x g, 5 min), pellet was resuspended in 4 ml complete medium and seeded in 65-mm Falcon culture dishes, pretreated with 0.1 mg/mL Poly-L-lysine. Each pup was processed separately. At sub-confluence (5–10 days in vitro; DIV), cells were detached with trypsin and plated for experiments. At this stage, microglial cells were removed by magnetic-activated cell sorting (MACS) negative selection using anti-CD11b-conjugated beads (Cat. 130-093-636) and MS magnetic columns (Cat. 130-042-201), (both from Miltenyi Biotec, Bologna, Italy). As controlled by anti-GFAP (astroglial marker), anti-Iba1 (microglial marker) and anti-MAP2 (neuronal marker, all from Merck Millipore, Billerica, MA) immunocytochemistry, resulting cultures contained virtually no microglial and neuronal cells (Ruffinatti et al., 2018).

Mixed astrocyte-neuron cultures were prepared as described previously (Grolla et al., 2013) with slight modifications. After enzymatic and mechanical dissociation, final cellular pellet was resuspended in neurobasal A medium (Invitrogen, Cat. 10888022) supplemented with 2% B27 supplement (Invitrogen, Cat. 17504044), 2 mg/mL glutamine, 10 U/mL penicillin and 100 mg/mL streptomycin, and plated as described above. Two/third of medium volume was changed every 5^th^ day and the cells were lysed at days in vitro (DIV) 12-15. For enriched neuronal cultures, cytosine β-D-arabinofuranoside (40 μM, Cat. C6645, Sigma) at DIV4 and neurons were lysed at DIV12.

### 2.5. Total RNA extraction and real-time PCR

Tissues or cells were lysed in Trizol reagent (Invitrogen, Cat. 15596026). Total RNA was extracted using Absolutely RNA miRNA kit (Agilent, 400809) according to manufacturer’s instructions. 0.5-1 μg of total RNA was retrotranscribed using random hexamers and ImProm-II RT system (Promega, Milan, Italy, Cat. A3800). Real-time PCR was performed using iTaq qPCR master mix according to manufacturer’s instructions (Bio-Rad, Segrate, Italy, Cat. 1725124) on a SFX96 Real-time system (Bio-Rad). To normalize raw real-time PCR data, S18 ribosomal subunit was used. Sequences of oligonucleotide primers is listed in Supplementary Table 1. The real-time PCR data are expressed as delta-C(t) of gene of interest to S18 allowing appreciation of the expression level of a single gene.

### 2.6. Lysates preparation and Western blotting

Astrocytes were grown in 100 mm dishes (1×10^6 /dish), while mixed neuron-astroglial cultures were grown in 65 mm dishes-(0.5 mln cells/dish). Cells were lysed (lysis buffer: 50 mM Tris HCl (pH 7.4), sodium dodecyl sulfate (SDS) 0.5%, 5 mM EDTA) and 20 μg proteins were resolved on 12% SDS-polyacrylamide gel. Proteins were transferred onto nitrocellulose membrane using Trans-Blot Turbo Transfer System (Bio-Rad), blocked in skim milk (5%, for 1 h; Cat. 70166; Sigma) and immunoblotted with indicated antibody. Rabbit anti-GAPDH (1:5000, Sigma Cat. G9545) or anti-β-actin (1:10000, Sigma, Cat. A1978) were used to normalize protein load. Primary antibodies were: rabbit anti-CaNB1 (1:500; Millipore Corp. Cat. 07-1494); mouse anti-PSD95, clone K28/43, antibody (1: 1000, Cat. MABN68, Merck Millipore, Vimodrone, Italy); mouse anti-Sy38 (1:500, Immunological Sciences, Cat. MAB-10321). Goat anti-mouse IgG (H+L) horseradish peroxidase-conjugated secondary antibody (1: 8000; Cat. 170-6516, Bio-Rad) was used. The protein bands were developed using SuperSignal West Pico chemiluminescent substrate (Cat. 34078; Thermo Scientific, Rockford, IL, USA). Densitometry analysis was performed using Quantity One software (Bio-Rad).

### 2.7. Electrophysiology

#### 2.7.1. Hippocampal slice preparations and CA1 pyramidal neurons electrophysiology

The mice were anesthetized with halothane (Sigma, St. Louis, MO) and killed by decapitation. Brains were carefully removed and placed in an ice-cold solution containing (mM): 87 NaCl, 2.5 KCl, 25 NaHCO_3_, 1.25 NaH_2_PO_4_, 75 Sucrose, 25 Glucose, 0.2 CaCl_2_, 7 MgCl_2_, bubbled with 95% O_2_ and 5% CO_2_. Brains were glued with cyanoacrylate glue to a support placed inside the cutting chamber of the vibratome (1000 plus, Vibratome, USA) and cut in 300 μm coronal slices. The slices were placed in Artificial cerebrospinal fluid aCSF solution containing (mM): 125 NaCl, 2.5 KCl, 1.25 NaH_2_PO_4_, 26 NaHCO_3_, 10 Glucose, 2 CaCl_2_, 1 MgCl_2_ for one hour at 34–35°C and thereafter for at least one hour at room temperature before recordings (Cunha et al., 2018). We performed whole-cell patch-clamp recordings from CA1 pyramidal neurons. Recordings were performed under continuous perfusion (1–1.5 ml min^−1^) with a bath solution containing (in mM): NaCl (119), KCl (5), HEPES (20), glucose (30), MgCl_2_ (2), CaCl_2_ (2) and in the same extracellular solution added with 10 μM Gabazine (SR95531) to block GABA-A receptors (Abcam) and maintained at 32°C. To record intrinsic properties of neurons, we used patch pipettes (resistance between 5.5–7 MΩ), filled with an intracellular solution consisting of (mM): 138 K-gluconate, 8 KCl, 10 Hepes, 0.5 EGTA, 4 Mg-ATP, 0.3 Na-GTP adjusted to pH 7.3 with KOH and 300 mOsm. The intrinsic excitability of CA1 was measured by using the current clamp mode of the whole-cell patch-clamp technique: CA1 resting membrane potential was set at −80 mV and the cells were then injected for 2 s or 800 ms with depolarizing current steps (50 pA in a 25 pA increment).

#### 2.7.2. Cerebellum Slice preparation and cerebellar granule cells electrophysiology

The mice were anesthetized with halothane (Sigma, St. Louis, MO) and killed by decapitation in order to remove the cerebellum for acute slice preparation according to a well-established technique (Armano et al., 2000; Sola et al., 2004). The cerebellar vermis was isolated and fixed on the vibroslicer’s stage (Leica VT1200S, etc) with cyano-acrylic glue. Acute 220 μm-thick slices were cut in the parasagittal plane in cold cutting solution containing (in mM): 130 potassium gluconate, 15 KCl, 0.2 EGTA, 20 Hepes and 10 glucose, pH adjusted to 7.4 with NaOH. Slices were incubated for at least 1 h at room temperature (24–25°C) in oxygenated extracellular Krebs solution containing (in mM): 120 NaCl, 2 KCl, 2 CaCl_2_, 1.2 MgSO_4_, 1.2 KH_2_PO_4_, 26 NaHCO_3_, 11 glucose, pH 7.4, when equilibrated with 95% O_2_ and 5% CO_2_. Slices were later transferred to a recording chamber mounted on the stage of an upright microscope (Zeiss, Oberkochen, Germany) and perfused (1–1.5 ml min^−1^) with the same extracellular solution added with 10 mM Gabazine (SR95531) to block GABA-A receptors (Abcam) and maintained at 32°C. Where indicated, ouabain (0.1 μM, Sigma) was added 5 min prior to recording. Patch pipettes were pulled from borosilicate glass capillaries (Hilgenberg, Malsfeld, Germany) and, when filled with the intracellular solution, had a resistance of 7–9 MΩ before seal formation. The whole-cell recording pipettes were filled with the following solution (in mM): 126 potassium gluconate, 4 NaCl, 5 Hepes, 15 glucose, 1 MgSO_4_.7H_2_O, 0.1 BAPTA-free, 0.05 BAPTA-Ca2+, 3 Mg_2_+-ATP, 0.1 Na+-GTP, pH 7.2 adjusted with KOH. This calcium buffer is estimated to maintain free calcium concentration around 100nM. We performed whole-cell patch-clamp recordings from CGCs in acute cerebellar slices. Recordings were performed with a Multiclamp 700-B amplifier (3dB; cutoff frequency (fc), 10 kHz), data were sampled with a Digidata 1440A interface, and analyzed off-line with pClamp10 software (Axon Instruments, Molecular Devices, USA). Membrane current and potential were recorded using the voltage-clamp mode and the current-clamp mode of the amplifier, respectively (D’Angelo et al., 1995). To examine the intrinsic excitability of CGCs we used the current clamp mode of the whole-cell patch-clamp technique: CGC resting membrane potential was set at −80 mV and the cells were then injected for 2 s or 800 ms with depolarizing current steps (from 8 to 48 pA in a 2 or 4 pA increment).

#### 2.7.3. Basal synaptic transmission

Evoked excitatory postsynaptic currents (EPSCs) were recorded stimulating the afferent fibers using a bipolar tungsten electrode (WPI) connected to a stimulator (npi electronics) through an isolator unit, delivering voltage pulses (10-15V) of 250 μs duration. Electrical stimulation was delivered to mossy fibers and Schaeffer collaterals in cerebellar and hippocampal slices, respectively. In order to isolate excitatory transmission, the GABA-A receptor blocker SR-95531 (100 μM) was added to the extracellular solution. For analysis, EPSC raw traces were low-pass filtered at 1 kHz and the peak amplitude was measured using pClamp 10 software (Axon Instruments, Molecular Devices, USA). The average amplitude is reported as mean ± SEM.

#### 2.7.4. Resting membrane potential of cultured astrocytes

Resting membrane potential (RMP) recordings were performed at room temperature in the whole-cell configuration using an Axopatch 200A (Axon Instruments). Cells were bathed in a solution containing (in mM): 138 NaCl, 4 KCl, 2 CaCl2, 1 MgCl2, 10 glucose and 10 HEPES at pH 7.25 adjusted with NaOH. Whole-cell patch pipettes were filled with (in mM): 140 KCl, 2 NaCl, 5 EGTA, 0.5 CaCl2 and 10 HEPES at pH 7.25 adjusted with KOH (Rocchio et al., 2019). Pipettes had a resistance of 3-5 MΩ. Soon after the patch rupture, membrane resting potential values were digitized with Digidata 1440A (Molecular Devices, USA) and recorded using pClamp software (Molecular Device, USA) at a sampling frequency of 1 kHz. A single measurement was obtained by averaging the first 30 s of the RMP trace.

### 2.8. Na^+^/K^+^ ATPase (NKA) assay

To measure NKA ATPase activity hippocampal or cerebellar tissue from one animal per point were used. Purified astroglial and enriched neuronal cultures were grown in 6 well plates at an approximate density of 1 × 10^5^ cells per well. Cell or tissues were lysed in an ATPase/GTPase activity assay kit buffer, and free phosphate was measured as according to manufacturer’s instructions (ATPase/GTPase activity assay kit, Cat. MAK113-1KT, Sigma). Ouabain (3 mM, Cat. 1076, Tocris) was added and NKA-specific ATPase activity was calculated as a difference between Ouabain and vehicle-treated samples. When whole-tissue lysates were processed, FK506 (200 nM) was added together with Ouabain. For determination of NKA ATPase activity in astroglial and neuronal cultures, FK506 was added to culture medium (200 nM) 24 h prior lysate preparation. Data are expressed as pmol of consumed inorganic phosphate per μg of total proteins per minute of reaction (pmol/μg/min).

### 2.9. Assessment of glutamate uptake and release

Primary astrocytes were cultured in 24-well plates at 5×10^4^ cells /well until 80% of confluence. For glutamate uptake measurement, cells were treated in DMEM/F12 medium with 5mM glutamate (L-glutamic acid monosodium salt, Sigma, Cat. No. G1626) with or without TBOA ((3S)-3-[[3-[[4-(trifluoromethyl)bemzoyl]amino]phenyl]methoxy]-L-aspartic acid, Tocris, Cat. No. 2532). After 90 min of incubation, supernatants were collected, centrifuged (16,000×g for 10min at 4°C) and analyzed for residual glutamate using the protocol described by the manufacturers (Amplex® Red Glutamic Acid/Glutamate Oxidase Kit, Invitrogen, Cat. No. A12221). Glutamate uptake was calculated as difference of fluorescence at 590 nm between glutamate plus TBOA and glutamate alone and normalized to protein concentrations of corresponding cell lysate.

For detection of glutamate release, cells were washed three times with 0.9% NaCl and left in culture in 0.9% NaCl plus 4% of sucrose. After 3h of incubation, supernatants were collected, centrifuged (16,000×g for 10min at 4°C) and analyzed for glutamate released using the protocol described by the manufacturers (Amplex® Red Glutamic Acid/Glutamate Oxidase Kit.

### 2.10. Statistical analysis

For statistical analysis and related graphical representations, GraphPad Prism v.7 and G*Power 3.1 (Faul et al., 2007) were used. Normally distributed data were expressed as mean ± SEM and parametric hypothesis tests have been used whenever possible, after evaluating the normality of residuals and the homoscedasticity of the dataset (Shapiro-Wilk and Levene’s test, respectively). In particular, unless otherwise specified, two-sample comparisons were performed using the unpaired two-tailed Student’s *t*-test, while, for comparisons between more than two experimental groups, ANOVA (one- or two-way, according to the number of independent variables) was used along with HSD Tukey post-hoc test. When the above assumptions were not met, parametric tests have been avoided in favor of specific nonparametric alternatives and median and interquartile range (IQR) were used as measures of central tendency and dispersion respectively. In all cases, tests were conducted at the significance level α=0.05 and the exact *p*-values are reported over the scatter-bars in the graphs. Additionally, for the most critical comparisons (namely when failing in rejecting the null hypothesis using a sample size smaller than n=6 per condition), a statistical power sensitivity analysis has been carried out to determine the minimum detectable effect (MDE) with α=0.05 and a statistical power π=0.8. Results of such a sensitivity analysis are reported in figure captions (Figs. 3A-B, 5E, 7B-C) in terms of both Cohen’s *d* and minimum detectable difference in dependent variable values (Δ), taking advantage of the estimated (pooled) standard deviation of the population(s).

## 3. RESULTS

### 3.1. Generation and validation of conditional astroglial CaN KO mouse line

A mouse model with conditional KO of CaN in GFAP-expressing astrocytes was generated by crossing calcineurin B1 (CaNB1)^flox/flox^ mice (Neilson et al., 2004) with 77.6 GFAP-Cre mice (Gregorian et al., 2009) resulting in CaNB1^flox/flox^:GFAP-Cre^+/−^ (<u>a</u>stroglial <u>c</u>alci<u>n</u>eurin conditional <u>k</u>nock-<u>o</u>ut, ACN-KO) and CaNB1^flox/flox^:GFAP-Cre^−/−^ (ACN-Ctr) mice (Suppl. Fig. 1). It is known that, in such brain regions as cerebellum (CB) and hippocampus (Hip), nearly all astrocytes are GFAP-positive (Verkhratsky and Nedergaard, 2018); hence, in these brain regions nearly all astrocytes will have deletion of CaN. We, therefore, used these two brain regions for validation and characterization of ACN-KO mice. At the mRNA level, primers that span the deleted region in the CaN gene (from the 2^nd^ to the 6^th^ exon), amplify only a 517 bp band in ACN-Ctr, while a shorter band of 95 bp appears in ACN-KO CB (Fig. 1A) and Hip (Suppl. Fig. 2A) due to elimination of exons 3-5 from the CaNB1 gene in a sub-population of cells, likely to be GFAP^+^ astrocytes. Notably, in PCRs using primers that amplify only the deleted RNA region (exons 3-5, Fig. 1B and Suppl. Fig. 2B) amplicons were present in whole tissue preparations from both ACN-KO and ACN-Ctr with no differences in band intensity, strongly suggesting that CaNB1 expression in neurons is unaltered. As expected, furthermore, Cre mRNA was present only in ACN-KO tissues (Fig. 1C and Suppl. Fig. 2C). To formally demonstrate astrocyte-specific CaNB1 deletion, we made use of fluorescence-activated cell sorting (FACS) and isolated Draq5^+^/GLAST^+^/CD90.2^−^ astrocytes from the CB of postnatal day 20 (P20) ACN-KO and ACN-Ctr mice (Suppl. Fig. 3). In these preparations, primers amplifying exons 3-5 failed to detect CaNB1 mRNA from FACS-purified astrocytes by real-time PCR (Fig. 1D). Abundant expression of CaN, including CaNB1, in neurons, instead, did not allow direct validation of CaNB1 KO in astrocytes by immunohistochemistry (Suppl. Fig. 4A). Nevertheless, a reduced background staining around intensely stained cerebellar Purkinje cell dendrites can be appreciated in ACN-KO as compared to ACN-Ctr tissues as revealed by measuring line scan profile and image intensity standard deviation in anti-CaNB1 stained sections (Suppl. Fig. 4B). At protein level, CaNB1 was strongly reduced in purified astroglial cultures from CB and Hip of ACN-KO mice (Fig. 1E,F) but it was not significantly changed in mixed neuron-astroglial cultures (Fig. 1G). To further demonstrate CaNB1 deletion in astroglial cells, we crossed ACN-KO mice with Rosa26-tdTomato reporter mice (kind donation from Prof. Frank Kirchhoff, University of Saarland, Homburg, Germany, (Jahn et al., 2018)). We enzymatically dissociated hippocampal tissues from P15 pups expressing tdTomato, plated dissociated cells, and performed immunocytochemical analysis of CaNB1 expression at 48 h after plating. We found a significantly lower expression of CaNB1 protein in ACN-KO compared with ACN-Ctr mice (Supp. Fig. 4C).

**Figure 1.**
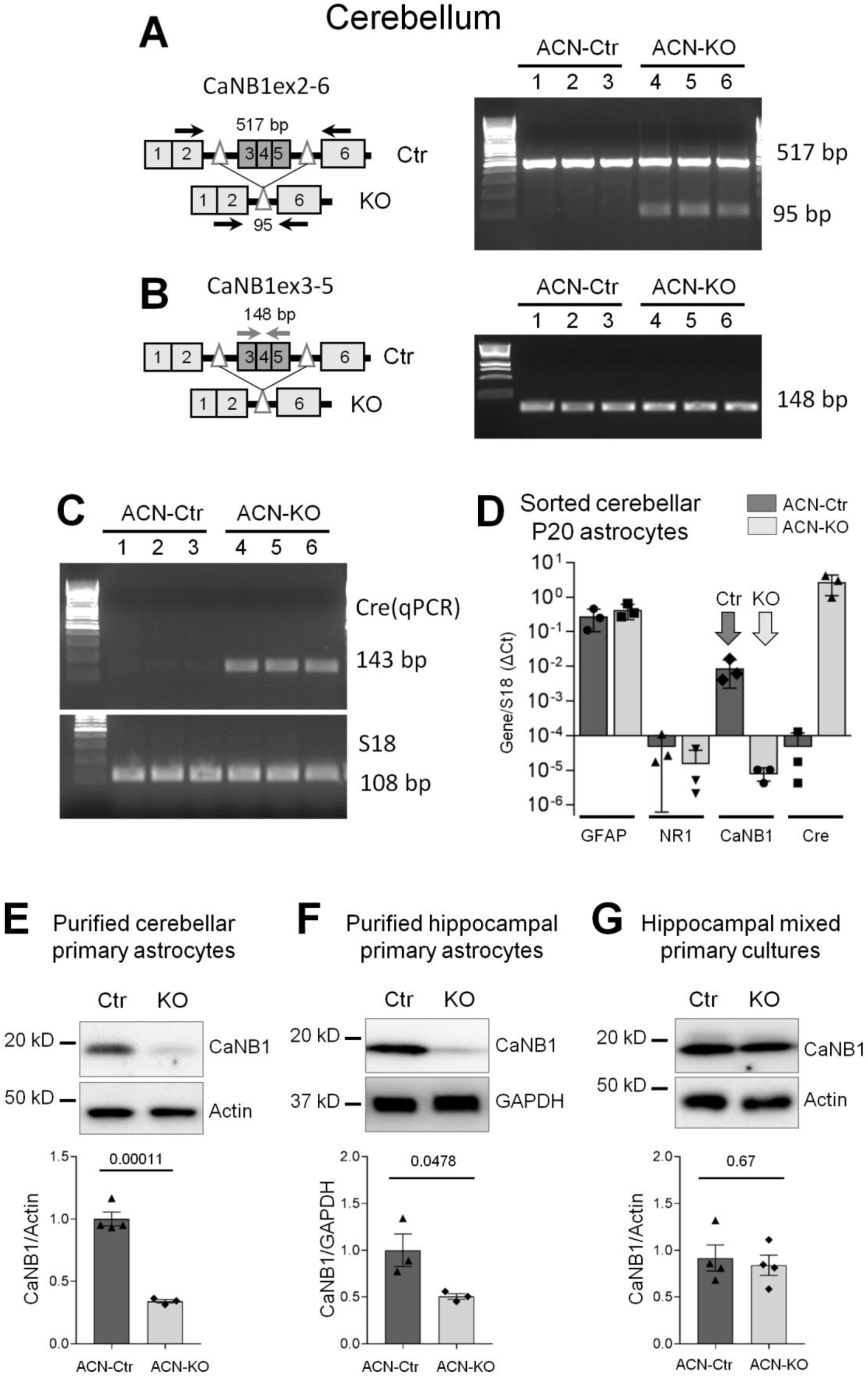
PCR characterization of ACN-KO mice. **A**, RT-PCR on whole cerebellar tissue using primers spanning exons 2-6 of CaNB1 mRNA. Alongside the 517 bp amplicon present in both WT and KO, the appearance of 95 bp amplicon in ACN-KO tissue indicates deletion of exons 3-5. Primers amplifying the deleted exons 3-5 (148 bp) show the presence of amplicons in both ACN-Ctr and ACN-KO tissues (**B**), while Cre amplicon (143 bp) was present only in ACN-KO cerebellum (**C**). Representative PCRs are shown from at least 4 independent litters. S18 mRNA (108 bp) was used as loading control. **D**, Real-time PCR on FACS sorted Draq5^+^/GLAST^+^/CD90.2^−^ astrocytes from cerebellum of P20 ACN-KO mice (mean ± SD, n = 3). Primers were used to amplify specifically exons 3-5 which are deleted in ACN-KO astrocytes (light-grey arrow). **C** and **D**, Anti-CaNB1 western blot analysis of purified cultures of cerebellar (**A**) and hippocampal (**B**) astrocytes shows strong reduction of CaNB1 protein in ACN-KO astrocytes while mixed neuron-astroglial primary cultures still retain calcineurin presence (**C**). Data are presented as mean ± SEM of 3-4 independent tissue preparations.

### 3.2. Postnatal onset of CaNB1 KO and normal neurogenesis, growth and development of ACN-KO mice

Recently, another CaNB1^flox/flox^/GFAP-Cre mouse line has been generated by using the human GFAP promoter. The Authors report defects in nutrition and premature death of pups around 1 mo of age (Fujita et al., 2018). Here we report that, in contrast to the mouse described by (Fujita et al., 2018), ACN-KO mice breed normally and deliver 6.6 ± 2.7 pups per litter, which is not different from that reported for the C57Bl/6 strain (Verley et al., 1967). No differences were observed in growth and body weight between ACN-Ctr and ACN-KO littermates (Fig. 2A). Likewise, there was no difference in brain weight between ACN-KO and wild type (WT) mice by 1 mo (Fig. 2B). There were no differences in the macrostructure of CB and Hip (Suppl. Fig. 5A) as well as in external appearance of ACN-KO and ACN-Ctr mice (Suppl. Fig. 5B).

**Figure 2.**
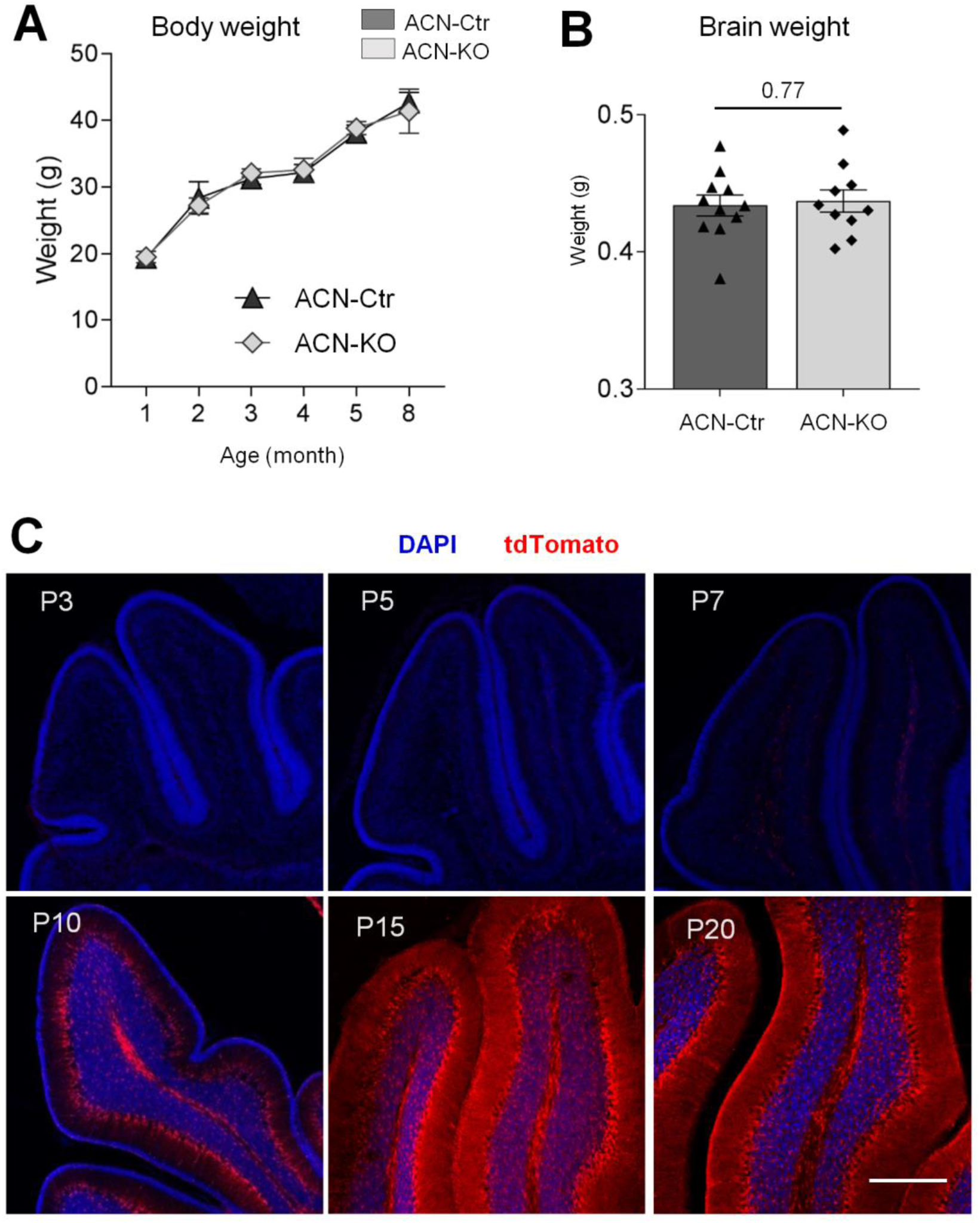
Normal growth and brain weight of ACN-KO mice, and time course of CaNB1 KO development in the cerebellum. Body weight until 8 mo of age (**A**) and brain weight at one month of age (**B**) were not different between ACN-Ctr and ACN-KO mice. In **A**, data are expressed as mean ± SEM. The number of mice per age was 6-20. In **B**, data are expressed as mean ± SEM of 11 (ACN-Ctr) and 10 (ACN-KO) brains. **C**, Time course of tdTomato expression in the cerebellum of RATO-ACN-KO mice shows appearance of reporter protein between 7 and 10 postnatal days. Full and widespread CaNB1 KO in the cerebellum is achieved by P15-P20. Images are representative from at least 3 ACN-KO mice for each age-point. Bar, 300 μm.

To investigate the spatio-temporal progression of CaNB1 KO we followed the development of tdTomato expression by confocal microscopy on fixed ACN-KO brain samples. The expression of the reporter protein was not evident before P7, increased steeply between P7 to P15 in both CB (Fig. 2C) and Hip (Suppl. Fig. 6A) and all astrocytes in these two brain regions expressed tdTomato at P20. To verify whether KO did not occur is some astrocytes, we analyzed the hippocampal formation of ACN-KO mice using the following strategy: taking into consideration that, in the Hip, astrocytes are organized in non-overlapping three-dimensional domains and that reporter tdTomato fills in the entire volume of astrocytes, we looked for the areas with no tdTomato expression, e.g., such “empty” areas are present in the cortex, in which not all astrocytes are GFAP-positive (Suppl. Fig. 7, insets A1 and B1). As shown in Suppl. Fig. 7, insets A2 and B2, at 1 mo of age, the entire volume of hippocampal formations was filled with tdTomato in both ACN-Ctr and ACN-KO, so that we were not able to detect boundaries between astrocytic domains. This indicates that 100% of astrocytes express tdTomato and therefore, achieved the full KO.

It has been stated that 77.6 GFAP-Cre line does not have Cre expression in neural stem cells in the Hip and other brain regions except the sub-ventricular zone (SVZ) (Gregorian et al., 2009). This is somewhat peculiar since a neural stem cell (NSC) marker Sox2 is typically co-expressed with GFAP during neurogenesis in perinatal period until the second postnatal week (Zhang and Jiao, 2015). It is therefore possible that there is an overlap between Sox2 and Cre expression in ACN-KO mice. In fact, we observed that a small proportion of Sox2 positive cells also expressed tdTomato reporter at P7 in both ACN-Ctr and ACN-KO (Suppl. Fig. 8A). If this would result in Cre expression in mature neurons, we could expect numerous neuronal cells expressing tdTomato reporter later in the life. Nevertheless, at 1 mo of age in the dentate gyrus there were only a few NeuN-positive neurons expressing tdTomato (Suppl. Fig. 8B), indicating a negligible effect of Cre expression in the neuronal progenitors on Cre occurrence in mature neurons.

Since adult neurogenesis impacts on neuronal excitability and functional plasticity (Lledo et al., 2006; Ikrar et al., 2013; Park et al., 2015), we quantified the number of migrating immature neurons (doublecortin (DCX) positive) and the number of newly generated neurons, (BrDU^+^/NeuN^+^ cells), in the subgranular (SGZ) and in the granular cell layer (GCL) within the dentate gyrus (DG) of the hippocampal formation. No differences were found in DCX^+^ or BrDU^+^/NeuN^+^ cells between ACN-Ctr and ACN-KO preparations (Fig. 3A,B). This demonstrates that adult hippocampal neurogenesis is not impaired by the deletion of CaNB1 in GFAP-expressing astrocytes, even if subtle effects cannot be excluded because of the limited statistical sensitivity of these comparisons (see figure legends). In addition, we evaluated whether levels of pre- and postsynaptic marker proteins were different between ACN-KO and ACN-Ctr mice. As shown in Fig. 3C and D, the amounts of synaptophysin and postsynaptic density protein 95 (PSD95) were not different in both CB and Hip tissues as well as in cultured primary hippocampal neurons.

**Figure 3.**
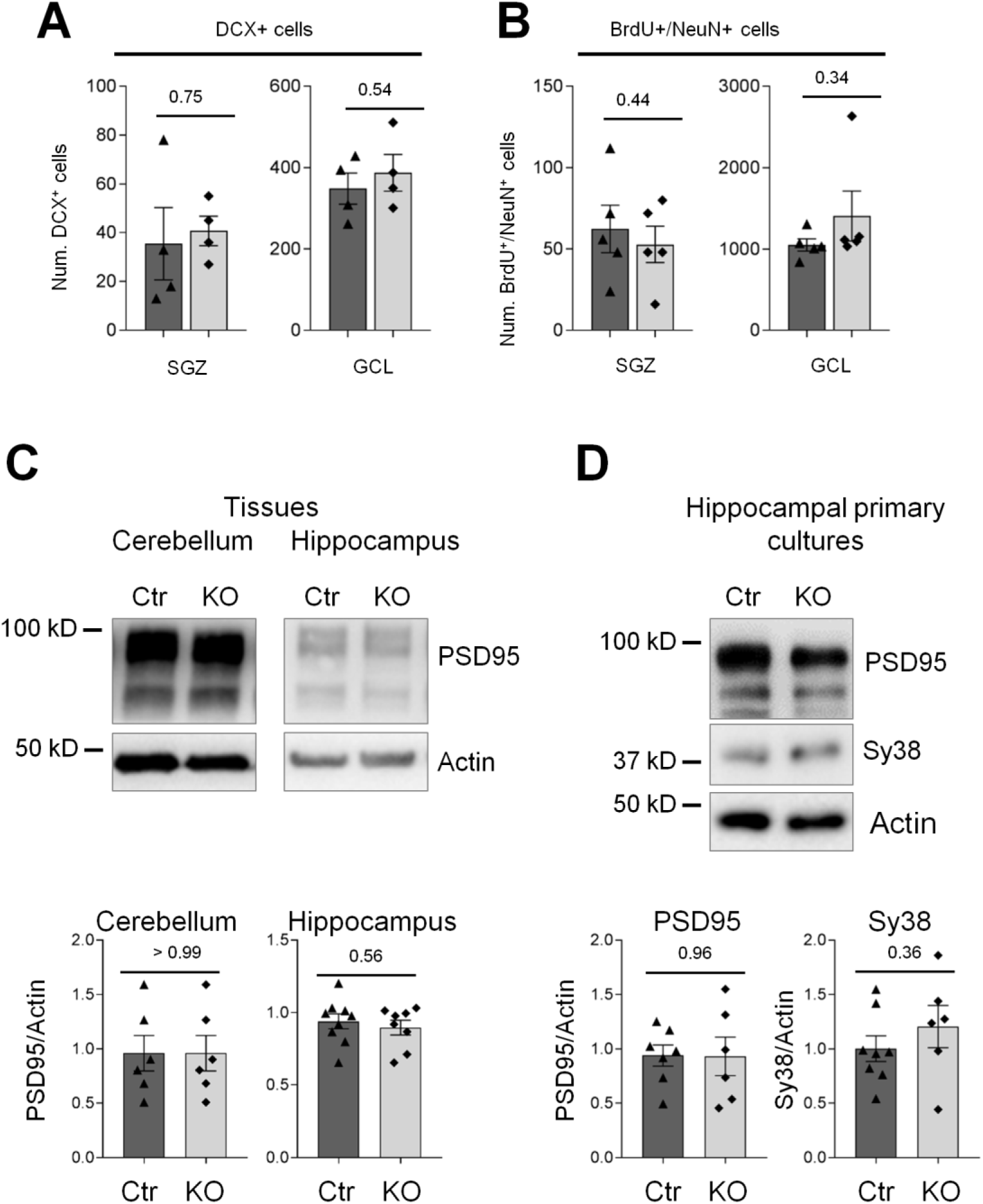
Adult neurogenesis and synaptic proteins are not altered in ACN-KO mice. **A**, Number of DCX^+^ cells was similar in ACN-Ctr compared to ACN-KO mice (n = 4 per group) in both subgranular zone (SGZ; MDE: *d* = 2.38, Δ = 47 cells) and granular cell layer (GCL; MDE: *d* = 2.38, Δ = 172 cells). **B**, Similarly, the number of BrdU^+^/NeuN^+^ cells was not significantly different between ACN-Ctr and -KO mice (n = 5 per group) in neither of these two brain regions (MDE in SGZ: *d* = 2.02, Δ = 53 cells; MDE in GCL: *d* = 2.19, Δ = 252 cells). Western blot analysis using anti PSD95 or anti-synaptophysin (Sy38) antibody revealed no differences in whole cerebella and hippocampal tissues (**C**), and in hippocampal mixed neuron-astroglial primary cultures (**D**). In **C** and **D**, data are expressed as mean ± SEM of 3-4 independent brain or culture preparations.

Taken together, we have generated an astrocyte-specific KO of CaNB1 without alterations in brain structure and development that is suitable to study the role of astroglial CaN in brain physiology.

### 3.3. Impairment of intrinsic excitability in hippocampal and cerebellar neurons in ACN-KO mice

Next, we proceeded to assess whether deletion of astroglial CaN affects neuronal intrinsic excitability in hippocampal CA1 pyramidal neurons and CGCs. Whole-cell patch-clamp recordings of hippocampal CA1 neurons (Fig. 4A) in response to current injection showed that ACN-KO neurons were unable to increase output spike frequencies following current injections to the levels shown by ACN-Ctr pyramidal cells (Fig. 4B), as shown by their input/output (I/O) relationship (Fig. 4C). In comparison to their Ctr counterparts, ACN-KO CA1 neurons demonstrated lower intrinsic excitability and generated less AP on current injection (Fig. 4C). Interestingly, no firing adaptation was observed in ACN-KO CA1 pyramidal cells, with no change in the after-hyperpolarization (AHP) evident during the discharge (Fig. 4B). However, voltage-dependent Na^+^ and K^+^ currents (Fig. 4D and Fig. 4E) and the passive membrane properties (Table 1) did not differ between ACN-Ctr and ACN-KO. These observations strongly suggest that the impairment of neuronal excitability in ACN-KO CA1 neurons depends on the dysregulation of extracellular K^+^ buffering, as described elsewhere (Meeks and Mennerick, 2004; Sibille et al., 2014; Chiacchiaretta et al., 2018).

**Figure 4.**
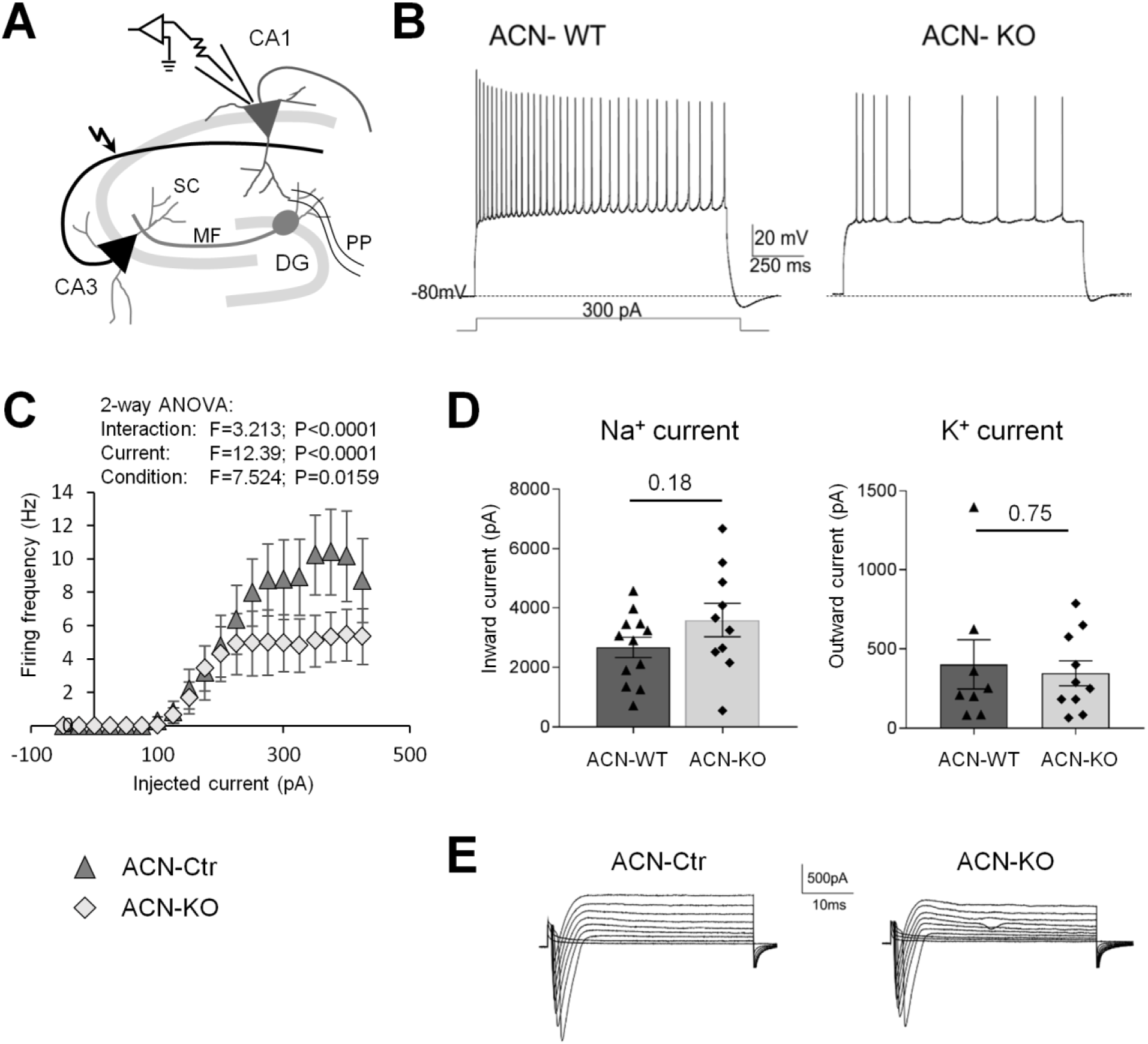
Evaluation of the intrinsic excitability of hippocampal CA1 neurons in ACN-Ctr and ACN-KO mice. **A**, Schematic representation of the hippocampal circuit. The basic electroresponsiveness of CA1 pyramidal neurons was investigated through the whole-cell patch-clamp technique. **B**, Electroresponsiveness of hippocampal CA1 pyramidal neurons. Voltage responses were elicited from −80 mV using step current injection. **C**, Relations between injected current and average firing frequency (F–I) for ACN-Ctr (n=9 slices from 5 mice) and ACN-KO (n=11 slices from 6 mice) CA1 pyramidal cells. Differences between the two F-I curves were statistically significant within the 325-400pA range, p=0.001. Data are reported as mean ± SEM. **D**, The bar charts compare inward (i.e. Na^+^) and outward (i.e. K^+^) current amplitudes measured at −40 mV and +20 mV in ACN-Ctr and ACN-KO mice. Data are reported as mean ± SEM. **E**. The traces show inward and outward currents elicited by increasing voltage steps from the holding potential of −70mV (capacitive transients have been truncated).

**Table 1.** Passive properties of cerebellar granule cells.

|  | ACN-Ctr (n=6) | ACN-KO (n=6) | p-value |
| --- | --- | --- | --- |
| **C<sub>m</sub> (pF)** | 2.67 ± 0.19 | 2.53 ± 0.13 | 0.58 |
| **R<sub>in</sub> (GΩ)** | 2.90 ± 0.30 | 3.06 ± 0.60 | 0.81 |
| **V<sub>rest</sub> (mV)** | -55.4 ± 3.34 | -47.2 ± 2.97 | 0.11 |

The input-output relationships of CGCs (Fig. 5A) revealed that the deletion of CaN from astrocytes shifted the high-frequency tonic firing into adaptive firing while depolarizing the cells (Fig. 5B). The I/O relationship (Fig. 4C) clearly showed that, at increasing depolarizing steps, the output firing frequency (22 pA: ACN-Ctr = 44.6 ± 15.5 Hz, n = 5; ACN-KO = 8.5 ± 3.74 Hz, n = 5; p = 0.05) was not able to follow the Ctr trend and soon dropped to a low number of spikes. In particular, ACN-KO CGCs showed rapid adaptation to high frequencies, associated to a decrease in the AHP depth of the spikes during the train (Fig. 5D). While Ctr neurons were able to maintain the AHP unvaried throughout the whole discharge, KO cells showed an average decrease in the AHP of about 25% of its initial value (p = 0.0312). Since the AHP phase of an AP is known to depend on K^+^ currents (D’Angelo et al., 1998), this effect might be due to a change in K^+^ currents or gradients in these cells. However, voltage-clamp current/voltage (I/V) relationship did not show any alteration in Na^+^ and K^+^ currents within the statistical sensitivity determined by the limited sample size (Fig. 5E and Fig. 5F). Moreover, the lower intrinsic excitability of ACN-KO granule cells was not due to the alteration of their passive properties (Table 2). Notably, the RMP showed a tendency towards more positive values in ACN-KO CGCs, although this difference was not statistically significant. Therefore, it is reasonable to speculate that the mechanisms of K^+^ gradient maintenance during high frequency activation are also impaired in ACN-KO CB. It is currently postulated that astrocytic Na^+^/K^+^ ATPase (NKA) maintains extracellular K^+^ homeostasis by clearing K^+^ released during neuronal firing thereby preventing the fall of the gradient across neuronal membrane which sustains intrinsic excitability (Larsen et al., 2016). We, therefore, hypothesized that astroglial CaN was able to modulate electro-responsiveness of CGCs and hippocampal CA1 neurons by regulating NKA activity and/or expression. If this is true, pharmacological blockade by ouabain is predicted to mimic the electrophysiological alteration recorded in ACN-KO mice. Therefore, we focused on CGCs, which, in addition to the remarkable adaptation of their firing frequency, show a large drop in the AHP depth during the discharge. Pre-incubating cerebellar slices with 0.1 μM ouabain, a concentration that selectively targets the astroglial alpha2 subunit (Clausen et al., 2017), converted the high-frequency tonic firing into an adaptive firing and reduced AHA (Fig. 5B–Fig. 5D), as observed in ACN-KO mice. Likewise, the passive properties were not affected by ouabain, although the positive shift in RMP was now significant (Table 3). This observation, therefore, strongly suggests that astroglial NKA expression and/or activity are compromised by deletion of astrocytic CaN.

**Figure 5.**
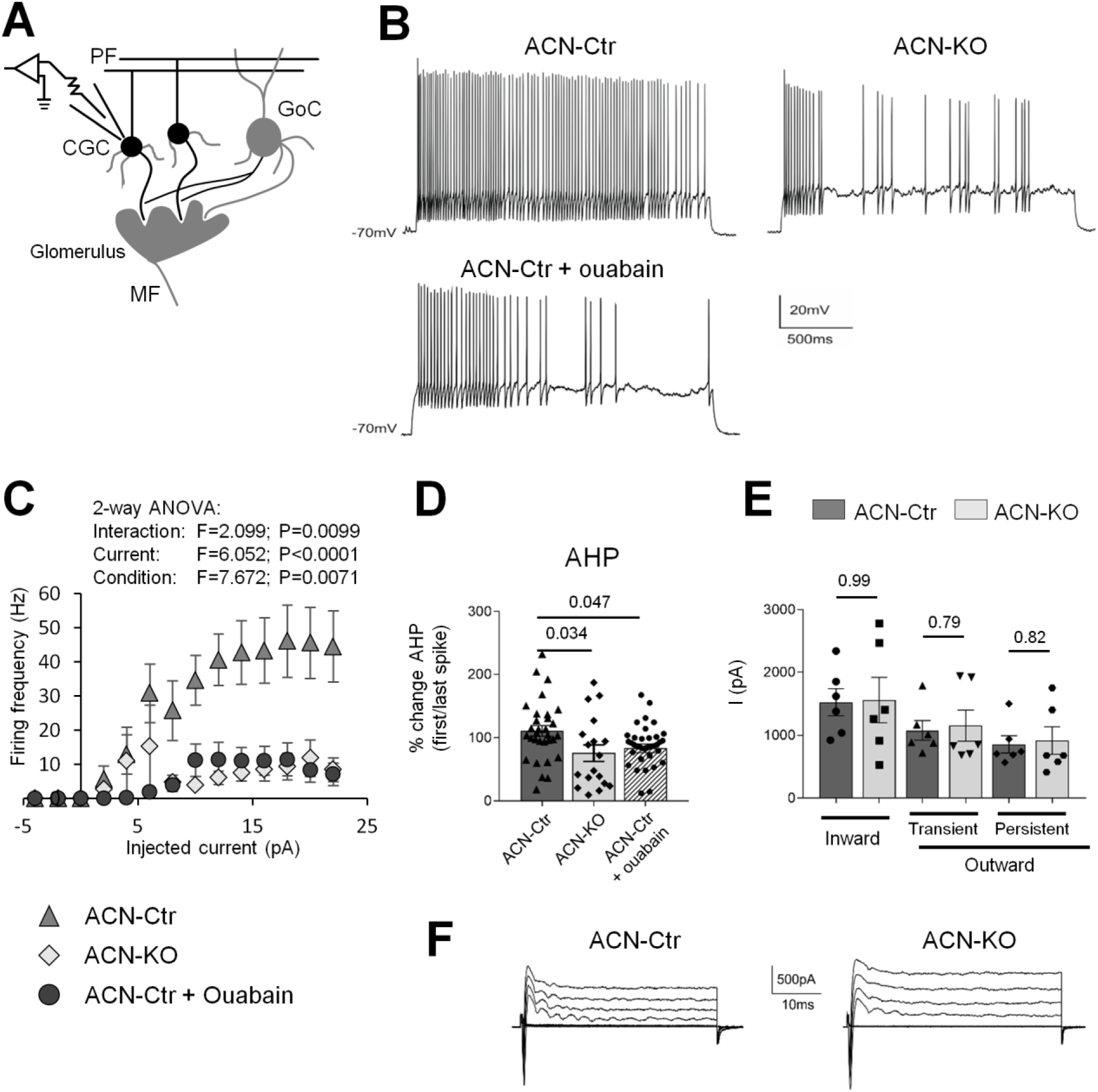
Evaluation of the intrinsic excitability of cerebellar granule cells in ACN-Ctr and ACN-KO mice. **A**, Schematic representation of a cerebellar circuit which illustrates how mossy fibers (MF) contact cerebellar granule cells (CGCs) and Golgi cell (GoC) dendrites. The basic electroresponsiveness of CGCs was investigated through the whole-cell patch-clamp technique. **B**, Granule cell electroresponsiveness. Voltage responses were elicited from −70 mV using step current injection. **C**, The plot shows the relationship between average firing frequency over 2 sec and the injected current intensity (F-I). n = 5 granule cells for all conditions (ACN-Ctr, ACN-KO and ACN-Ctr + ouabain (0.1 μM)). Data are reported as mean ± SEM. **D**, The bar chart shows the % change of AHP between the first and the last spike for both ACN-Ctr (n = 31 from 6 slices obtained from 6 different mice) and ACN-KO (n = 18 from 6 slices obtained from 6 different mice). **E**, The bar chart compares voltage-gated inward (i.e. Na^+^) and outward (i.e. K^+^) current amplitudes measured at −40 mV and +20 mV in ACN-Ctr (n = 6) and ACN-KO (n = 6) mice. No significant difference was detected in none of the performed comparisons. MDE for inward Na^+^ currents: *d* = 1.80, Δ = 1195.7 pA. MDE for transient outward K^+^ currents: *d* = 1.80, Δ = 828.4 pA. MDE for persistent outward K^+^ currents: *d* = 1.80, Δ = 731.5 pA. **F**, The traces show inward and outward currents elicited by increasing voltage steps from the holding potential of −70mV (capacitive transients have been truncated).

**Table 2.** Passive properties of hippocampal CA1 pyramidal cells.

|  | ACN-Ctr (n=4) | ACN-KO (n=4) | p-value |
| --- | --- | --- | --- |
| **C<sub>m</sub> (pF)** | 126.81±8.08 | 102.18 ±5.00 | 0.77 |
| **R<sub>in</sub> (GΩ)** | 0.222±0.03 | 0.256±0.008 | 0.10 |
| **V<sub>rest</sub> (mV)** | -52.2±4.24 | -55.7 ± 2.26 | 0.43 |

**Table 3.**
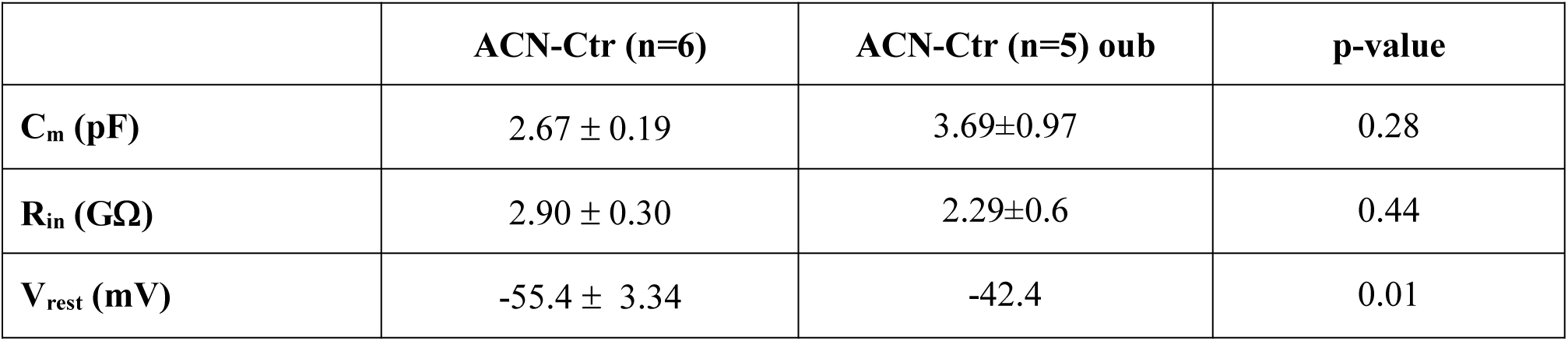
Passive properties of cerebellar granule cells upon preincuabation with 0.1 μM ouabain (15 min)

### 3.4. CaN-dependent impairment of NKA activity in astrocytes of ACN-KO mice

In order to address whether and how astroglial CaN regulates NKA, we first measured ouabain-sensitive ATPase activity in CB and Hip lysates as well as in primary hippocampal astrocytes from ACN-KO mice. Specific NKA activity was significantly inhibited in both CB and Hip whole lysates by 62% and 78%, respectively (Fig. 6A and Fig. B). Furthermore, NKA activity was inhibited by 67% in purified astroglial (Fig. 6C), but not neuronal (Fig. 6D), cultures from ACN-KO mouse hippocampi. Interestingly, in line with previous publications (Lea et al., 1994), NKA activity was inhibited by the CaN inhibitor FK506 in both total cerebellar lysates (Fig. 6A). However, when cell cultures were treated with FK506, NKA inhibition was observed only in astroglial cultures, but not in enriched neuronal cultures (Fig. 6C and D), which suggests that NKA requires the phosphatase activity of CaN in astrocytes but not in neurons.

**Figure 6.**
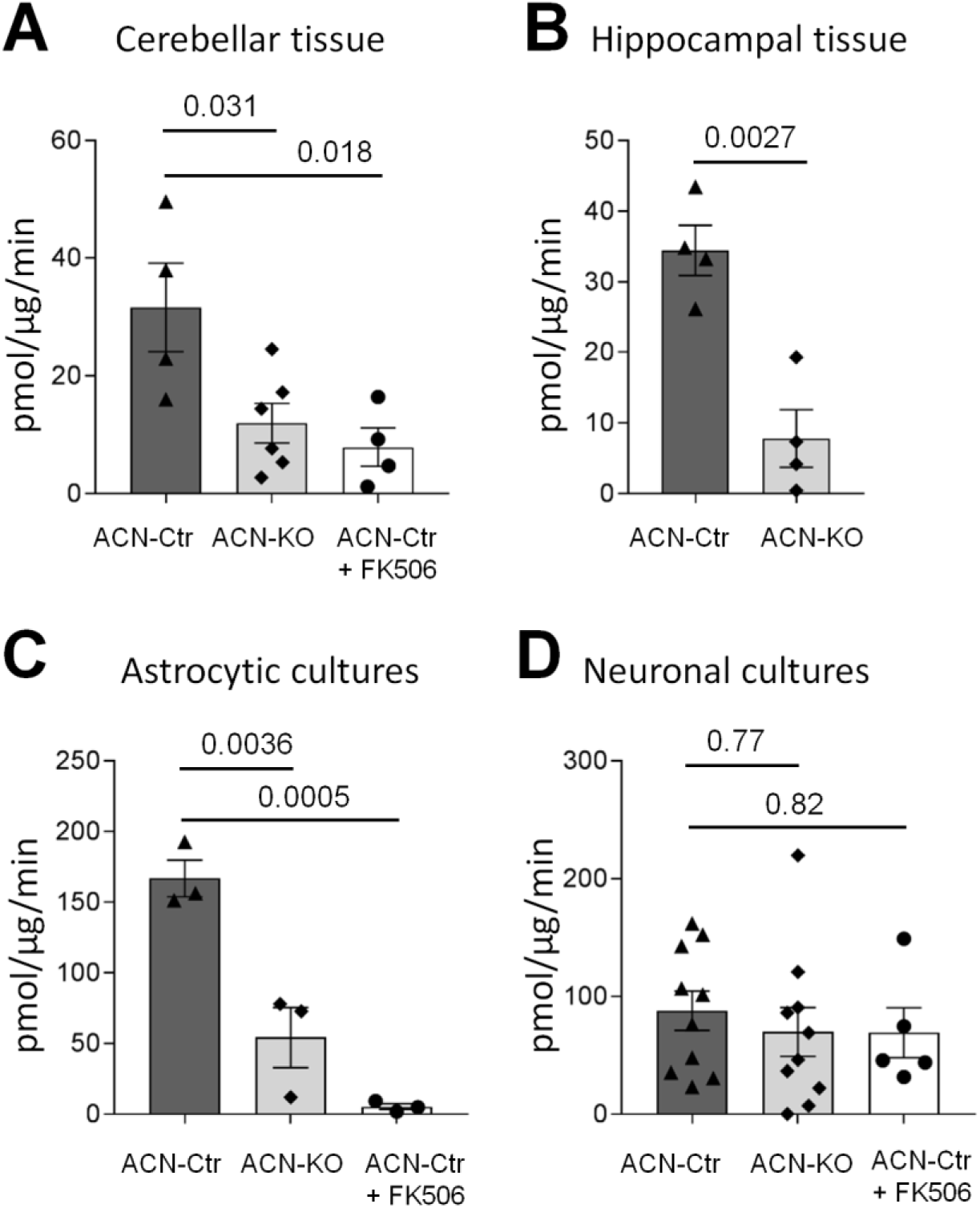
NKA activity is decreased in tissues and in cultured astrocytes of ACN-KO mice. Ouabain-sensitive ATPase activity was measured in whole tissue cerebellar (**A**) and hippocampal (**B**) lysates as well as in purified astroglial primary cultures (**C**) and in enriched neuronal cultures (**D**) from ACN-Ctr and ACN-KO mice. Where indicated FK506 was added to the lysates at a final concentration of 200 nM. NKA ATPase activity is expressed as pmol of inorganic phosphate per μg total proteins measured in 1 minute (pmol/μg/min). Data are expressed as mean ± SEM from 3-5 independent tissue or culture preparations.

Alongside direct dephosphorylation, CaN is known to regulate transcriptional activity of a number of genes in physiological as well as in pathological conditions. Transcriptional regulation may occur through direct activation of transcription factor of activated T-cells (NFAT) and by indirect activation of nuclear factor kappa-light-chain-enhancer of activated B cells (NF-kB) in astrocytes (Lim et al., 2013, 2016; Furman and Norris, 2014) and in other cell types (Hogan et al., 2003). NKA is composed of a larger catalytic alpha subunit and two smaller regulatory subunits (beta and gamma subunit) (Clausen et al., 2017). The functional core of NKA is composed of one alpha and one beta subunit, while the gamma subunit is required for fine tuning of NKA activity in tissue and isoform-specific manner (Geering, 2008). The alpha subunit includes four isoforms (coded by genes Atp1a1-4), of which three are expressed in brain (Atp1a1-3), the beta subunit includes three isoforms (coded by genes Atp1b1-3), of which two are expressed in brain. The heterodimer composed of alpha2 and beta2 is thought to be the astrocyte-specific variant, while the presence of alpha1 has also been shown (Larsen et al., 2016). We performed *in silico* analysis and found that the Atp1a2 displays four high affinity NFAT binding sites, two of which are located in the proximal promoter region (−203 bp and −264 bp) and two locating in the distal promoter region (−1843 bp and −1998 bp) upstream of transcriptional start site (Suppl. Fig. 9). Thus, a possible transcriptional effect of CaN on NKA expression and activity cannot be *a priori* ruled-out. Therefore, we employed real-time PCR to measure relative abundance of transcripts for Atp1a1 and Atp1a2 and Atp1b2 isoforms. We found, however, that during the entire period of brain development, specifically, at P0, P5, P10, P15, P20 and P30, in spite of strong developmental up-regulation (Orlowski and Lingrel, 1988) the levels of mRNA were not different between ACN-Ctr and ACN-KO (Fig. 7A). In line with this, protein levels of alpha1 and alpha2 NKA isoforms, as detected by WB, did not differ significantly either in CB or in Hip (Fig. 7B,C). Notably, despite their limited statistical sensitivity, these comparisons allow to rule out biologically relevant effects (fold changes greater than 2, see figure caption). Last, no differences were found in the expression and localization of the Atp1a2 NKA subunit in ACN-Ctr and ACN-KO cerebella by immunohistochemical analysis (Suppl. Fig. 12). These latter results suggest that activity modulation of NKA and not transcriptional regulation by CaN is important to control excitability of CB and Hip neurons by astrocytes.

**Figure 7.**
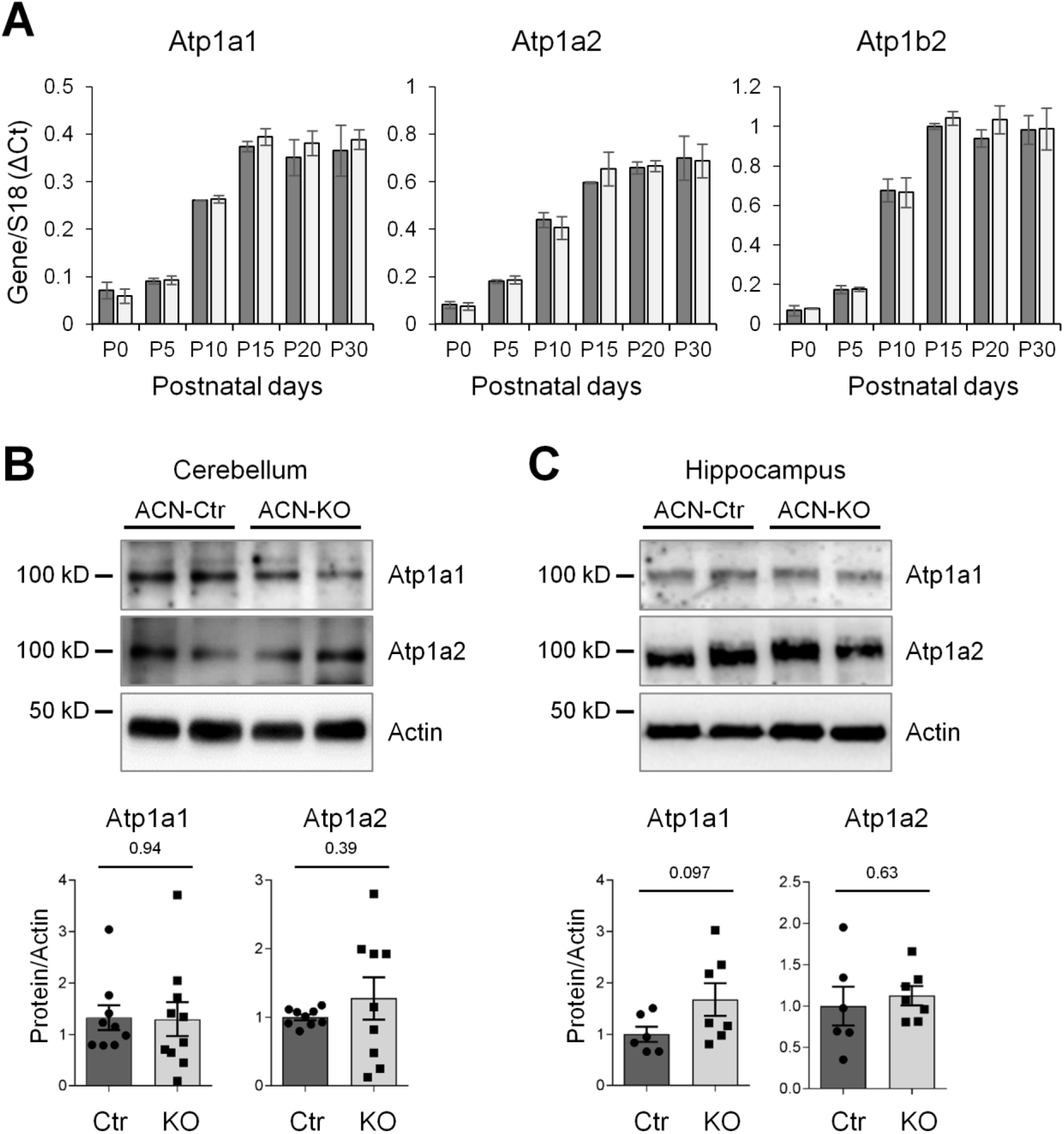
Transcripts and protein levels of astrocytic NKA isoforms are not different between ACN-Ctr and ACN-KO mice. mRNA levels of catalytic Atp1a1, Atp1a2 and regulatory Atp1b2 isoforms were measured by real-time PCR in hippocampal whole tissue lysates at indicated time points during the first month of post-natal development (**A**). Data are expressed as mean ± SEM of 3-6 hippocampal lysates. At each time-point, the differences were not significant between ACN-Ctr and ACN-KO. **B**, Western blot analysis could not detected any significant differential expression in whole tissue lysates from ACN-Ctr mouse cerebellum compared to ACN-KO neither for Atp1a1 (MDE: *d* = 1.40, Δ = 1.20) nor for Atp1a2 catalytic isoforms (MDE: *d* = 1.41, Δ = 0.88). **C**, Similarly, the differential expression of these two catalytic isoforms was not significant when evaluated in hippocampus (MDE for Atp1a1: *d* = 1.71, Δ = 1.05; MDE for Atp1a2: *d* = 1.71, Δ = 0.73). Representative bands are shown of 6-10 independent experiments.

Astrocytes may buffer extracellular K^+^ released during high frequency firing also through inwardly rectifying K^+^ channels (K_ir_) (Hertz and Chen, 2016). Indeed, the inwardly rectifying K_+_ channel 4.1 (K_ir_4.1) is specifically expressed in astrocytes and may be involved in K^+^ and maintenance of K^+^ permeability in resting conditions. In our conditions, both protein and mRNA K_ir_4.1 expression levels were slightly, although significantly, down-regulated in ACN-KO astrocytes (Suppl. Fig. 11A,B).

Last, using whole-cell current-clamp recordings, we have also investigated whether pharmacological inhibition of CaN may affect RMP in cultured astrocytes. The electrophysiological recordings revealed a range of RMPs from −89.7 to −23.8 mV (*n* = 16), with a median value of −70.7 mV and an interquartile range IQR = [−76.1; −57.1] mV. This wide range of RMPs is in agreement with previous results obtained in rat hippocampal slice astrocytes (McKhann et al., 1997). Then, we examined the effect of the inhibition of CaN activity on RMP. Overnight incubation of astrocytes with FK506 (200 nM) did not affect the median RMP, which resulted equal to −74.2 mV with an IQR = [−78.1; −66.5] mV (*n* = 16; *p*-value = 0.49, Mann-Whitney *U* Test, two-tail). Moreover, the marked heterogeneity of RMPs persisted in this condition, probably as an effect of a RMP bimodal distribution (McKhann et al., 1997; Bolton et al., 2006). In light of this, we focused our attention to the cellular subpopulation with a RMP more negative than −55 mV, which also represents the majority of the cell sample. Again, we found no significant difference between control and treated astrocytes (mean ± SEM in control astrocytes was −75.2 ± 2.4 mV (*n* = 12); in treated astrocytes was −76.3 ± 1.3 mV (*n* = 12; *p*-value = 0.69, *t*-test, two-tail). These observations confirm that astroglial CaN deletion specifically affects neuronal firing by modulating astroglial NKA activity.

### 3.5. Na^+^ homeostasis, glutamate uptake and gliotransmitter release in ACN-KO cultured astrocytes

The findings reported so far clearly show that NKA activity is impaired in ACN-KO mouse astrocytes and regulates neuronal excitability, but not astroglial RMP. Nevertheless, the impairment of astroglial NKA activity could affect astrocytes by altering their intracellular Na^+^ homeostasis and interfering with Na^+^-dependent mechanisms, such as Na^+^-dependent glutamate transporters (Kirischuk et al., 2012; Verkhratsky et al., 2013). We, therefore, made use of primary hippocampal cultures as well as primary purified astrocytes from ACN-Ctr and ACN-KO mice to investigate the link between astroglial CaN, NKA activity and Na^+^-dependent astroglial functions. First, using the Na^+^-sensitive fluorescent dye SBFI, we found that the intracellular Na^+^ concentrations were significantly higher in ACN-KO (31.62 ± 2.36 mM, n = 279) than in ACN-Ctr astrocytes (9.1 ± 1.01 mM, n = 343, p < 0.0001) (Fig. 8A). This observation suggests that the Na^+^-dependent transport may be altered in ACN-KO astrocytes. However, the glutamate uptake rate was not different between ACN-KO (19.85 ± 1.65 nmol/μl/min) and ACN-Ctr cells (21.58 ± 3.45 nmol/μl/min, p = 0.66) (Fig. 8B). This may be explained by differences in the glutamate transporters expression. We primed a recently published microarray database (Ruffinatti et al., 2018) to evaluate the glutamate transporter primarily expressed in cultured astrocytes and found that glutamate/aspartate transporter (GLAST) is predominantly expressed over glutamate transporter 1 (GLT-1) (Fig. 8D). Thus, we investigated GLAST expression using WB analysis. Fig. 8E shows that GLAST is expressed at significantly higher levels in ACN-KO than in ACN-Ctr astrocytes, which may compensate the intracellular Na^+^ increase and the decrease in Na^+^ driving force for glutamate uptake. Last, we measured release of gliotransmitter glutamate and found that it was significantly lower in ACN-KO (15.39 ± 0.42 nmol/μl/min) than in ACN-Ctr astrocytes (19.19 ± 1.56 nmol/μl/min) (Fig. 8C). Nevertheless, when we investigated basal excitatory transmission, we did not find any difference in ACN-Ctr and ACN-KO excitatory postsynaptic currents (EPSCs), elicited by electrical stimulation of hippocampal Schaeffer collaterals or cerebellar mossy fibers, as illustrated in Supplementary Figure 9.

**Figure 8.**
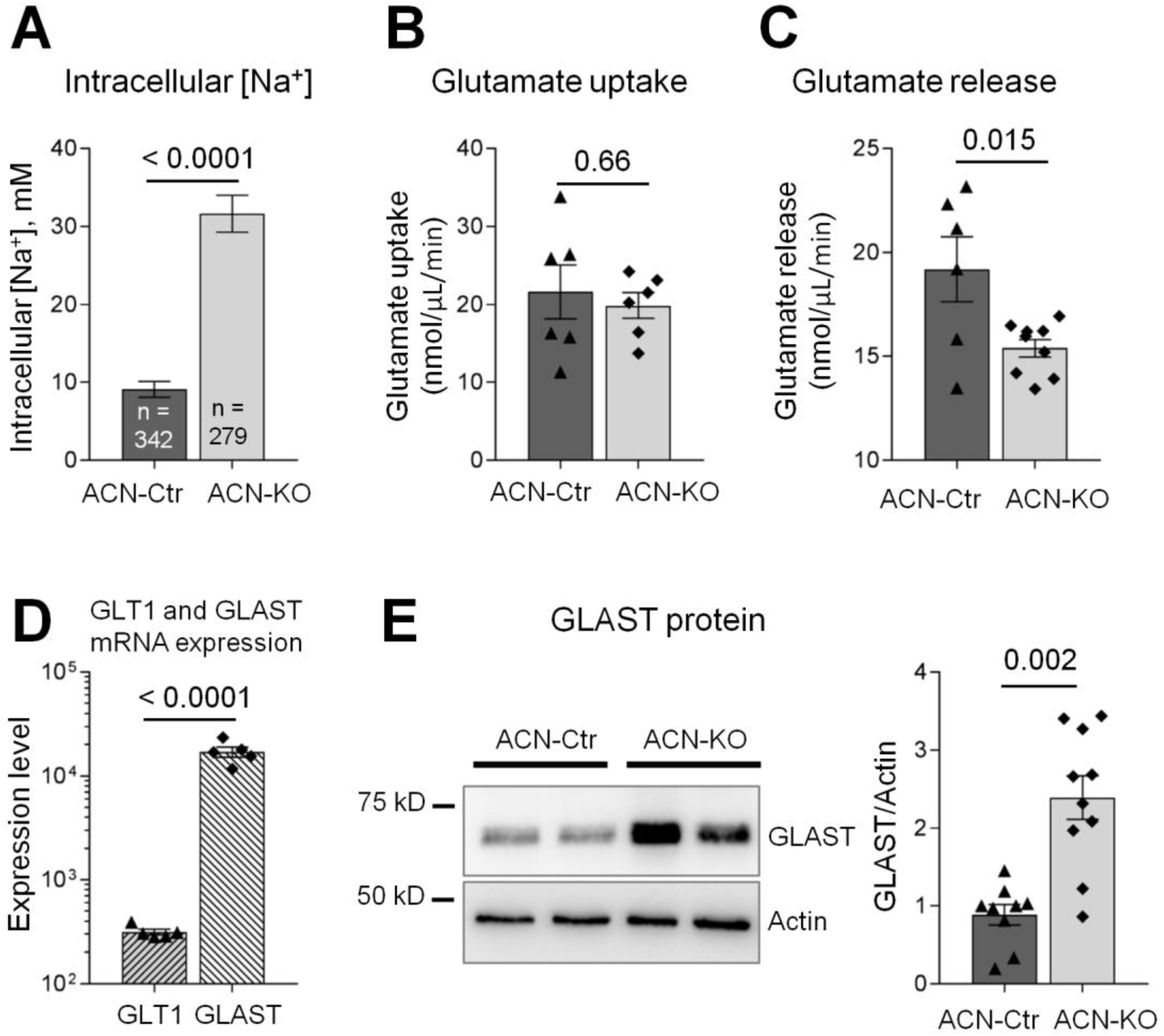
Intracellular Na^+^ and Na^+^-dependent transport in ACN-KO asctrocytes. **A**, intracellular [Na^+^] was measured using SBFI probe in cultured hippocampal astrocytes. Data are expressed as mean ± SEM of n = 343 cell (ACN-Ctr) and n = 279 cells (ACN-KO) from 5 independent cultures for each genotype performed in triplicate. Glutamate uptake (**B**) and release (**C**) was assessed in 6-9 independent culture preparations. **D**, to compare relative expression of GLT1 and GLAST mRNA, a previously published microarray database obtained on cultured hippocampal astrocytes was primed (Ruffinatti et al., 2018). Data are expressed as mean ± SEM fluorescence levels of 5 independently prepared astroglial cultures. **E**, GLAST protein expression in primary cultures of hippocampal astrocytes from ACN-Ctr and ACN-KO mice. Data are expressed as mean ± SEM 9 and 10 independent cultures for ACN-Ctr and ACN-KO, respectively.

Finally, we have investigated the reactive status of astrocytes in ACN-KO mice. Immunostaining with anti-GFAP antibody showed comparable results in the hippocampal formation of both ACN-Ctr and ACN-KO mice (Suppl. Fig. 13A). No differences were found in the number of GFAP-positive cells and in the area of GFAP staining (Suppl. Fig. 13B and C). We next, employed real-time PCR to assess expression levels of astroglial marker GFAP, microglial reactivity marker Iba1 and inflammation-related cytokines IL-1β and TNFα. No significant differences were found in mRNA levels of Iba1, IL-1β and TNFα (Suppl. Fig. 14).

Collectively, these data suggest that there may be alterations in Na^+^-regulated processes in ACN-KO astrocytes, although the functional consequences of such alterations remain to be fully investigated.

## 4. DISCUSSION

In the present manuscript, we investigated for the first time the role of astrocytic CaN in the healthy brain. We report that the specific deletion of CaNB1 from GFAP-expressing astrocytes during the postnatal period: 1) does not alter development of ACN-KO mice and of the morphology of the central nervous system (CNS); 2) does not have impact on the number of hippocampal NPCs; 3) severely impairs intrinsic excitability in granule cerebellar neurons and hippocampal CA1 pyramidal neurons; and 4) that this the impairment is likely due to the lack of positive regulation of NKA by calcineurin in astroglia.

To generate the ACN-KO mouse, we used a 77.6 GFAP-Cre mouse line recently reported and deposited to Jackson Lab mouse repository by Michael Sofroniew (JaxLab Stock No. 024098). To date, several GFAP-Cre lines are commercially available, which differ in cellular specificity and developmental pattern of Cre expression (Zhuo et al., 2001; Garcia et al., 2004; Gregorian et al., 2009). Recently, Fujita and colleagues (Fujita et al., 2018) reported the generation of a CaN KO using Cre expression driven by the a 2.2 kb 5’ flanking fragment of the human GFAP gene generated by (Zhuo et al., 2001). The authors observed alterations of early postnatal development and premature death of pups around 1 mo of age. The authors concluded that the deletion of CaNB1 from glial cells of the enteric nervous system resulted in strong inflammation and mucosal permeability in small intestine which led to an inability to digest and absorb food (Fujita et al., 2018). In preliminary experiments in our lab, we observe expression of tdTomato reporter also in enteric glial cells, thereby confirming this localization and the ablation of calcineuin in these cells also in our model. Yet, we do not confirm the effect on growth and development. It has to be noted that the GFAP-Cre line used by Fujita et al. displays embryonic (E13.5) onset of Cre expression and in adulthood is also expressed in oligodendroglia, ependymal cells, some neurons in the CNS as well as in periportal liver cells (Zhuo et al., 2001) introducing thus a bias in the interpretation of the results. For example, neuronal CaN plays a crucial role in axonal growth and guidance during vertebrate development by activating NFAT (Nguyen and Di Giovanni, 2008). Also the expression in pre-natal periods might explain the difference. For example, a recent study demonstrated that early embryonic exposure to cyclosporine, another established CaN inhibitor, causes a reduction in eye and brain size in zebrafish (Clift et al., 2015). Although it has been stated that the 77.6 GFAP-Cre mouse line does not present Cre expression in neuronal progenitors in the Hip (Gregorian et al., 2009), we found partial co-localization of a NSC marker, Sox2, with tdTomato reporter at P7. This finding suggests a short temporal overlap between the end of the embryonic and perinatal Sox2 expression and the onset of GFAP-Cre expression in ACN-KO mice. Nevertheless, this did not result in Cre expression in adult neurons, as at 1 mo of age we have found only single cells in the DG which were positive for both NeuN, a marker of differentiated neurons, and tdTomato reporter. Altogether, these data suggest that the ACN-KO mouse presented in this work is useful for investigating the role of astroglial CaN in physiology and pathology of CNS.

A central mechanistic finding of this report is that selective deletion of astrocytic CaNB1 severely impaired intrinsic excitability in CGCs and hippocampal CA1 neurons. It has long been known that alterations of astrocyte capacity to clear extracellular K^+^ result in dramatic changes in neuronal network excitability, which may result in neurological disorders, including epilepsy, Rett syndrome, Alzheimer’s Disease and Huntington’s Disease (Meeks and Mennerick, 2004; Sibille et al., 2014; Chiacchiaretta et al., 2018). Originally, two mechanisms were proposed for astrocytes to buffer extracellular K^+^: (i) the net K^+^ uptake, an energy-dependent K^+^ buffering mediated by glial NKA (Hertz, 1965, 1979) and (ii) the K^+^ spatial buffering, an energy-independent re-distribution of K^+^ from regions with high K^+^ to those with low K^+^ concentrations (Orkand et al., 1966). Subsequent work showed that the K^+^ spatial buffering involved K_ir_ channels for local K^+^ accumulation and gap junctions for redistribution of K^+^ (Hertz and Chen, 2016). Currently, it is postulated that astrocytic K_ir_-mediated spatial buffering is mainly responsible for resting K^+^ permeability (Neusch et al., 2001; Djukic et al., 2007; Kucheryavykh et al., 2007), while K^+^ clearance during neuronal firing is mainly due to NKA activity. Notably, analysis of K^+^ affinity of heterodimers with different alpha and beta subunits composition revealed that the astroglial alpha2-beta2 combination has the lowest K^+^ affinity (Larsen et al., 2014) and therefore is particularly suitable for the removal of large amounts of K^+^ released from neurons during intense neuronal activity (Larsen et al., 2016). The following evidences strongly support the hypothesis that the inhibition of astrocytic NKA impairs neuronal excitability in ACN-KO mice. First, NKA activity was strongly inhibited in hippocampal and cerebellar tissues and in purified astroglial cultures, but not in enriched neuronal cultures from ACN-KO pups. This finding demonstrates that astrocytic NKA is selectively inhibited in the brain of ACN-KO mice. Second, NKA activity was significantly impaired by FK506 in both total cerebellar lysates and in cultured astrocytes from Ctr animals. This finding demonstrates that CaN activation is permissive towards astrocytic NKA activity. This latter observation supports reports by others in cells other than astrocytes (Marcaida et al., 1996; Mallick et al., 2000; Woolcock and Specht, 2006; Rambo et al., 2012) as well as in kidney tissue preparations (Tumlin and Sands, 1993; Lea et al., 1994). It is postulated that CaN-dependent dephosphorylation activates, while phosphorylation, *e.g.* by protein kinase A (PKA) or protein kinase C (PKC), inhibits NKA activity (Bertorello et al., 1991; Bertuccio et al., 2003, 2007; Poulsen et al., 2010). Third, the lack of significant changes in neuronal RMP indicates that deletion of astrocytic CaN did not alter the transmembrane K^+^ gradient in cerebellar and hippocampal neurons, which further confirms that neuronal NKA activity was unaltered. Finally, the pharmacological blockade with a concentration of ouabain that selectively targets the astroglial alpha2 subunit phenocopied the alterations in high frequency firing observed in ACN-KO CGCs. The slight, but statistically significant, positive shift in RMP could be explained by considering that 0.1 μM ouabain could also affect the neuronal alpha3 subunit of the NKA, which is expressed in the granular layer of the CB (Bøttger et al., 2011), thereby directly depolarizing CGCs. Of note, although the alteration in astroglial Na^+^ homeostasis observed in ACN-KO astrocytes resulted in a decrease in glutamate release, basal excitatory synaptic transmission was not altered either in Hyp or in CB. Future work, however, will have to confirm these data *in vivo* and to assess whether short- and long-term synaptic plasticity is impaired in ACN-KO mice. Likewise, the functional implication of K_ir_4.1 down-regulation will have to be further assessed. Accordingly, K_ir_4.1 down-regulation is predicted to depolarize the RMP, thereby increasing neuronal excitability (Tong et al., 2014), which is conversely reduced in ACN-KO mice while there is no significant alteration in the RMP of either hippocampal CA1 neurons or CGCs.

It should be acknowledged that only the astrocyte-specific knock-down or knock-out of NKA would unequivocally rule-out the involvement of neuronal NKA and other K^+^ transporters, e.g. K_ir_ channels, in the observed inhibition of neuronal excitability in ACN-KO mice. However, such experiments might be challenging because, at worse, complete elimination of NKA ATPase activity from astrocytes would require knocking down of at least two NKA isoforms, namely alpha1 and alpha2. This rationale is based on an assumption that both alpha1 and alpha2 NKA are inhibited by deletion of CaNB1 in ACN-KO astrocytes. Even if the lower K^+^ affinity of alpha2-beta2-composed NKA (Larsen et al., 2014, 2016) suggests the predominant involvement of alpha2 isoform in K^+^ clearance, the contribution of alpha1 subunit cannot presently be ruled-out. Moreover, cultured purified astrocytes tend to lose the specific alpha2 isoform, thereby making the alpha1 the predominant isoform (Peng et al., 1998). For these considerations, we did not investigate which NKA isoform is activated by astrocytic CaN, acknowledging this as a limitation of the present work.

In addition to regulation of target proteins by dephosphorylation, CaN regulates protein expression by modulating gene transcription, e.g. through direct interaction with transcription factor NFAT in many cell types (Hogan et al., 2003), including astrocytes (Furman and Norris, 2014). The presence of several high score NFAT-binding sites in the promoter of Atp1a2 gene, detected by our *in silico* analysis, suggests that both of these mechanisms may potentially concur for the control of NKA activity in astrocytes. The hypothesis of transcriptional regulation of NKA by CaN is also supported by extensive data on the yeast analog of NKA, ENA1, in which stress-induced expression of ENA1 is mediated by CaN through activation of the transcription factor Crz1 (Mendoza et al., 1994; Mendizabal et al., 2001; Ruiz et al., 2003; Ruiz and Ariño, 2007). Nevertheless, our data demonstrate that, in the current experimental setting, mRNA levels of both Atp1a1 and Atp1a2 were not different throughout the postnatal brain development period, a finding which rules-out transcriptional regulation. In agreement with this assertion, protein levels of both alpha1 and alpha2 NKA were not different in hippocampal and cerebellar tissues of ACN-Ctr and ACN-KO mice.

The regulation of neuronal excitability by astrocytic CaN reinforces the emerging evidence in favor of bidirectional communication between neurons and astrocytes (Durkee and Araque, 2019). Recently, we have proposed that neuronal activity may lead to CaN activation in non-neuronal cells (Lim et al., 2016) as it was shown for other cells, *e.g.* in pericytes (Filosa et al., 2007) and in brain microvascular endothelial cells (Zuccolo et al., 2017). Furthermore, we have previously demonstrated that chemical induction of neuronal activity robustly activates CaN in astrocytes juxtaposed to activated neurons in *in vitro* co-cultures (Lim et al., 2018). Based on pharmacological characterizations, we have suggested that these effects are mediated by glutamate through astroglial mGluR5 and store-operated Ca^2+^ entry mechanism (Lim et al., 2018).

In summary, we present here a conditional KO of CaN in astrocytes suitable for investigating the role of CaN in the CNS physiology. Initial characterization of the mouse model led to establish that this enzyme in astrocytes regulates NKA activity and that neuronal excitability is altered both in the CB and in the Hip. From these data, a hypothetical model can be generated in which neurons signal to astrocytes and this leads to astrocytic Ca^2+^-rises that are sufficient to activate calcineurin, which in turn controls K^+^ transport and feeds back to neuronal excitability. More generally, our data show that astrocytes are key regulators of neuronal activity.

## Acknowledgements

This work was funded by grant 2014-1094 from the Fondazione Cariplo and grant “Bando Ricerca Locale” 2016 from The Università del Piemonte Orientale to DL; the Italian Association for Cancer Research (AIRC) and the Giovanni Armenise-Harvard Foundation to ADR; the Italian Ministry of Education, University and Research (MIUR): Dipartimenti di Eccellenza Program (2018–2022) - Dept. of Biology and Biotechnology "L. Spallanzani", University of Pavia, to FM; LT was supported by fellowship from the CRT Foundation (1393-2017); TS, LM and ED received funding from the European Union’s Horizon 2020 Framework Programme for Research and Innovation under the Framework Partnership Agreement No. 650003 (HBP FPA); TS and ED received founding from Centro Fermi (Roma, Italy) for the project Microcircuiti Neuronali Locali (MNL).

