## Supplemental Material for "Deletion of calcineurin from GFAP-expressing astrocytes impairs excitability of cerebellar and hippocampal neurons through astroglial Na^+^/K^+^ ATPase"

### Index

| Content | Page |
|---|---|
| Supplementary Methods | 2 |
| Supplementary Table 1. **List of oligonucleotide primers used for PCR** | 3 |
| Supplementary Figure 1. **Breeding scheme for generation of ACN-WT and KO mice** | 5 |
| Supplementary Figure 2. **Validation of CB1 KO in hippocampus by PCR** | 6 |
| Supplementary Figure 3. **FACS separation of GLAST$^{+}$ astrocytes** | 7 |
| Supplementary Figure 4. **Hist-cytological analysis of CB1 KO** | 8 |
| Supplementary Figure 5. **Normal brain development and appearance of ACN-KO mice** | 9 |
| Supplementary Figure 6. **Time course of CB1 KO in GFAP-expressing astrocytes** | 10 |
| Supplementary Figure 6. **Time course of CB1 KO in GFAP-expressing astrocytes** | 10 |
| Supplementary Figure 7. **Widespread Cre expression in ACN-KO mice** | 11 |
| Supplementary Figure 8. **Cre expression does not affect neural progenitors mature neurons** | 12 |
| Supplementary Figure 9. **Basal synaptic transmission is not altered in ACN-KO mice** | 13 |
| Supplementary Figure 10. **In silico analysis of NFAT-binding sites in Atp1a2 promoter** | 14 |
| Supplementary Figure 11. **Kir4.1 protein is downregulated in the ACN-KO hippocampus** | 15 |
| Supplementary Figure 12. **Histochemistry of Atp1a2 NKA subunit** | 16 |
| Supplementary Figure 13. **GFAP expression in the hippocampus of ACN-KO mice** | 17 |
| Supplementary Figure 14. **Inflammatory markers are not changed in ACN-KO mice** | 18 |

### SUPPLEMENTARY METHODS

#### In silico NFAT-binding site analysis.

For searching NFAT-binding sites in Atp1a2 promoter, a 3000 bp 5' untranslated region of Atp1a2 gene (Gene ID: 98660) was analysed by two transcription factor analyzing softwares: PROMO software Version 3.0.2 (http://alggen.lsi.upc.es/cgi-bin/promo_v3/promo/promoinit.cgi?dirDB=TF_8.3) using TRANSFAC database and GPminer (http://gpminer.mbc.nctu.edu.tw/index.php). High score NFAT-binding sites found by both search engines resulted in a list of 4 sequences, two in proximal promoter (-203 bp and -264 bp) and two in distal promoter region (-1843 bp and -1998 bp).

#### Genotyping protocol

For genotyping, small ear punch biopsies were taken and subjected to genomic DNA extraction using PCRBIO Rapid Extract PCR kit (Genzano di Roma (RM), Italy). Eight µL of crude DNA extract was used for each PCR reaction.

##### *GfapCre genotyping*

GfapCre was genotyped by real-time PCR using following forward (TCC ATA AAG GCC CTG ACA TC) and reverse (TGC GAA CCT CAT CAC TCG T) primers for GfapCre (JaxLab primers 15831 and 15832, respectively) and following forward (CAA ATG TTG CTT GTC TGG TG) and reverse (GTC AGT CGA GTG CAC AGT TT) primers for internal positive control (JaxLab primers oIMR8744 and oIMR8745, respectively). Real-time PCR was performed using iTaq qPCR master mix according to manufacturer's instructions (Bio-Rad, Segrate, Italy) on a SFX96 Real-time system (Bio-Rad). Amplification parameters were: (5 min @ 95°C) → 40 cycles of (20 sec @ 95°C – 30 sec @ 60.5°C – 20 sec @ 72°C).

##### *$CB1^{flox/flox}$ and RATO genotyping*

$CB1^{flox/flox}$ was genotyped using following forward (TTC GAG GAC AGC TAT ACA GAG AAA) and reverse (GAC CTC CAG CCT CCA CAT AC) primers (JaxLab primers 13938 and 13939, respectively). Expected band size is: 300 bp for $CB1^{flox/flox}$; 203 bp and 300 bp for $CB1^{flox/wt}$; and 03 bp for $CB1^{wt/wt}$.

RATO was genotyped using following primers: forward (AAGGGAGCTGCAGTGGAGTA), reverse (CCGAAAATCTGTGGGAAGTC) for RATO-WT and forward (GGCATTAAAGCAGCGTATCC) and reverse (CTGTTCCTGTACGGCATGG) for RATO-MUT (JaxLab primers).

PCR for both $CB1^{flox/flox}$ and RATO was performed using DreamTaq Green PCR Master Mix (Thermo Fisher) on Bio-Rad C1000 thermo cycler using following amplification protocol (5 min @ 95°C) → 34 cycles of (30 sec @ 95°C – 30 sec @ 62°C – 1 min @ 72°C) → (5 min @ 72°C). Amplified bands were resolved on 2.5 agarose gel and visualized using ethidium bromide.

**Supplementary Table 1. List of oligonucleotide primers used in this work.**

| Gene | Accession number | Forward Reverse | Sequence 5' to 3' |
|---|---|---|---|
| S18 | NM_213557 | Forward<br>Reverse | TGCGAGTACTCAACACCAACA<br>CTGCTTTCCTCAACACCACA |
| CaNB1ex2-6 | NM_024459.2 | Forward<br>Reverse | AGATGGGAAATGAGGCGAGT<br>AGTCACACATCCACCACCAT |
| CaNB1ex3-5 | NM_024459.2 | Forward<br>Reverse | GGCAACGGAGAAGTGGACT<br>CCTGGAAGAGTTCTCCATTGG |
| CaNB1qPCR | NM_024459.2 | Forward<br>Reverse | AGATGGGAAATGAGGCGAGT<br>TCCACGCTCAAAGAACCAGA |
| GFAP | NM_001131020.1 | Forward<br>Reverse | GCTCCAAGATGAAACCAACC<br>GAACTGGATCTCCTCCTCCA |
| GLAST | NM_148938.3 | Forward<br>Reverse | AATGCCTTCGTTCTGCTCAC<br>ATCCTCATGAGAAGCTCCCC |
| GLT1 | NM_001077514.3 | Forward<br>Reverse | CTGGTGCAAGCCTGTTTCC<br>TAGTTTCTTCAGGGGCCTCG |
| GS | NM_008131.4 | Forward<br>Reverse | ACTGTGAGCCCAAGTGTGT<br>AGGTACATGTCGCTGTTGGA |
| Kir4.1 | NM_001039484.1 | Forward<br>Reverse | CACTTCACCTTCGAGCCAAG<br>CCATTCTCACATTGCTCCGG |
| AQP4 | NM_001317729.1 | Forward<br>Reverse | GGTTGGAGGATTGGGAGTCA<br>GTTTGGAATCACAGCTGGCA |
| Aldh1l1 | NM_027406.1 | Forward<br>Reverse | GACGTGCTTCCAGATGACAC<br>CAATCAGTCGCACAGCCTG |
| Gabra1 | NM_010250.5 | Forward<br>Reverse | CATTCTGAGCACACTGTCGG<br>TGTCATAACCGTCCAGCAGT |
| Syt1 | NM_001252341.1 | Forward<br>Reverse | TGGGGAAGGGAAGGAAGATG<br>GCAGGTCACGACTAGAAGGA |
| KCC2 | NM_020333.2 | Forward<br>Reverse | CTGACGGACTGCGAGGAC<br>CCATGTTCCTGCCATCGTAC |
| NR1 | NM_008169 | Forward<br>Reverse | GCCAGGAGGAGAGACAGAGA<br>CATCACTCATTGTGGGCTTG |
| Atp1a1 | NM_144900.2 | Forward<br>Reverse | TGAGCTCAAGAAGGAAGTGTCT<br>GGATCTCAGCGGCCCTTG |
| Atp1a2 | NM_178405.3 | Forward<br>Reverse | CGATGAGCTGAAGAAGGAGG<br>CGTTGGGTCCATCTCTAGCC |
| Atp1b2 | NM_013415.5 | Forward<br>Reverse | GTTGAGGAGTGGAAGGAGTTC<br>GCTGAACATGGCCGTGAG |
| Iba1 | NM_019467.2 | Forward<br>Reverse | CCGTCCAAACTTGAAGCCTT<br>ACCCCAAGTTTCTCCAGCAT |
| IL-1β | NM_008361.3 | Forward<br>Reverse | AAGTTGACGGACCCCAAAAGA<br>TGTTGATGTGCTGCTGCGA |

| | | | |
|---|---|---|---|
| TNFα | NM_013693.2 | Forward<br>Reverse | ACTGAACTTCGGGGTGATCG<br>CTCCTCCACTTGGTGGTTTG |

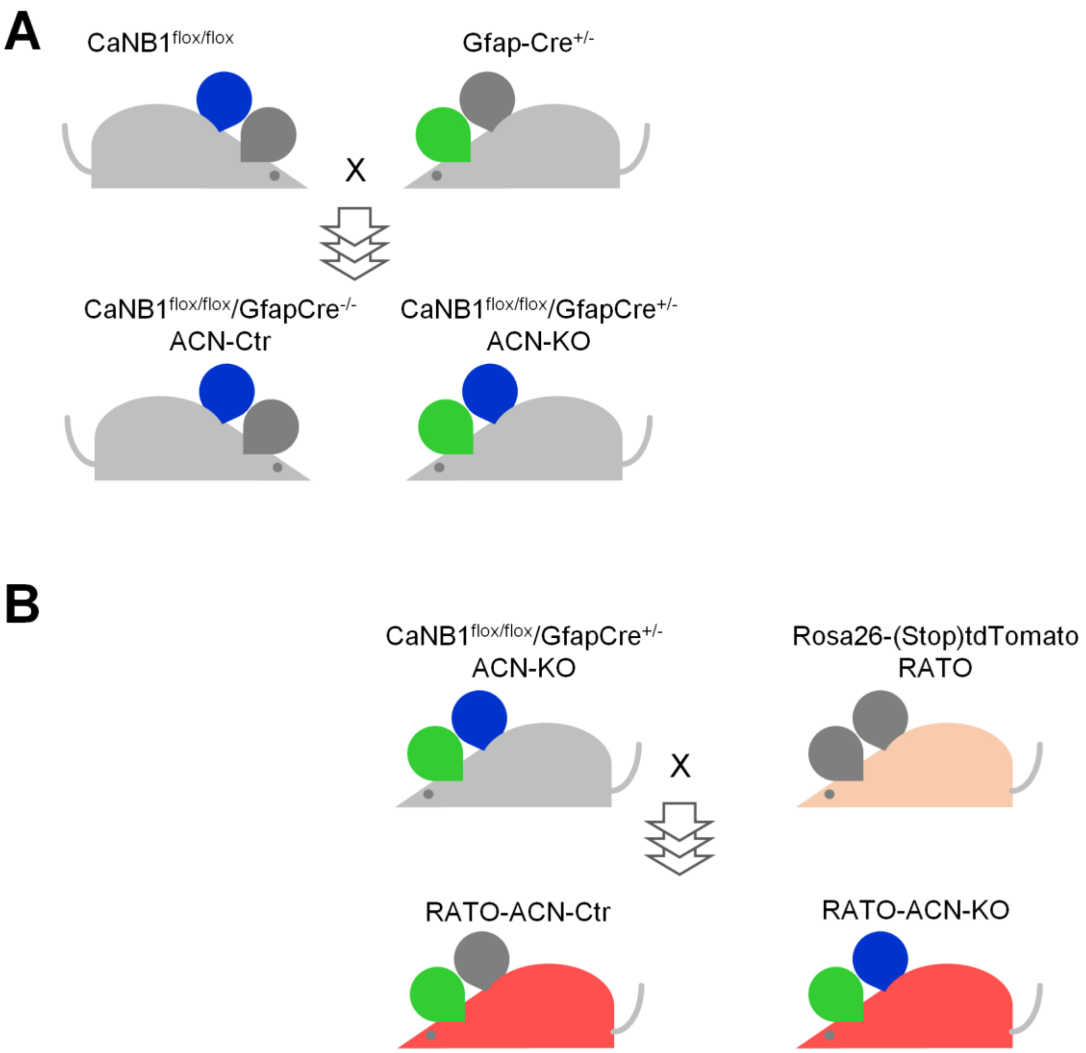


**Supplementary Figure 1. Breeding scheme for generation of ACN-Ctr and KO and reporter RATO-ACN-KO mice. A**, To generate CB1$^{flox/flox}$/GfapCre$^{-/-}$ (ACN-Ctr) and CB1$^{flox/flox}$/GfapCre$^{+/-}$ (ACN-KO) mouse lines (black frame), CB1$^{flox/flox}$ (Jax Lab strain B6;129S-Ppp3r1tm2Grc/J, stock number 017692) was crossed with Gfap-Cre$^{+/-}$ (Jax Lab strain B6.Cg-Tg(Gfap-cre)77.6Mvs/2J, stock number 024098). F1 Founders were backcrossed for at least 4 generations. **B**, To generate RATO-ACN-KO reporter mice, ACN-KO was crossed with Rosa26-CAG-flox-stop-flox-tdTomato (RATO) resulting in selection of RATO$^{flox/flox}$/ CB1$^{wt/wt}$/GfapCre$^{+/-}$ mice (RATO-ACN-Ctr) and RATO$^{flox/flox}$/ CB1$^{flox/flox}$/GfapCre$^{+/-}$ mice (RATO-ACN-KO). Blue ear, CB1$^{flox/flox}$; green ear, GfapCre$^{+/-}$; red head, RATO$^{flox/flox}$/GfapCre$^{+/-}$.

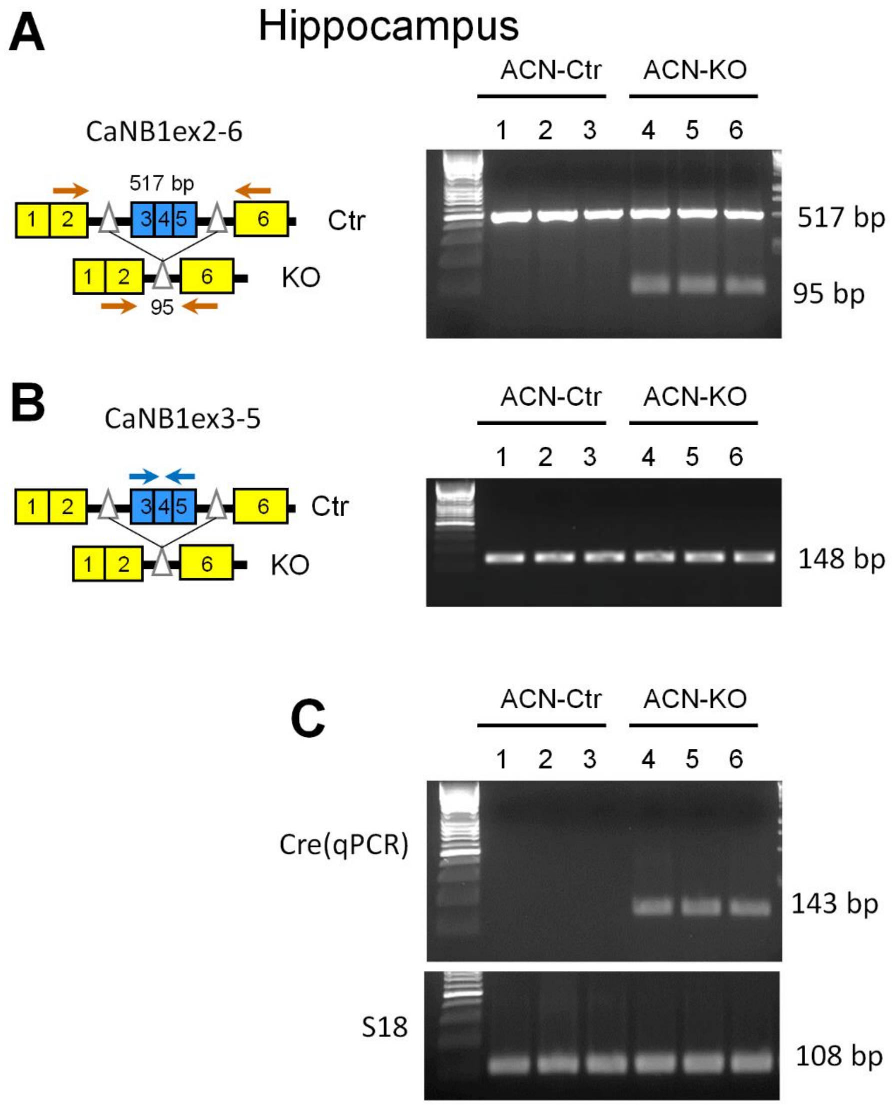


**Supplementary Figure 2. Validation of CaNB1 KO in the hippocampus of ACN-KO mice.** Primers design strategy for amplification of Exons 2-6 fragment (**A**) and Exons 3-5 (**B**). RT-PCR of whole hippocampal using Exons2-6 and Exons3-5 spanning primers shows appearance of 517 bp and 148 bp in both ACN-Ctr (lines 1-3) and ACN-KO (lines 4-6) tissues at one month of age (**A**). Note appearance of 95 bp band in Exons2-6 amplicon only in ACN-KO tissues. mRNA for Cre (143 bp) was amplified only in ACN-KO tissues (**C**). S18 mRNA amplicon (108 bp) was used as loading control.

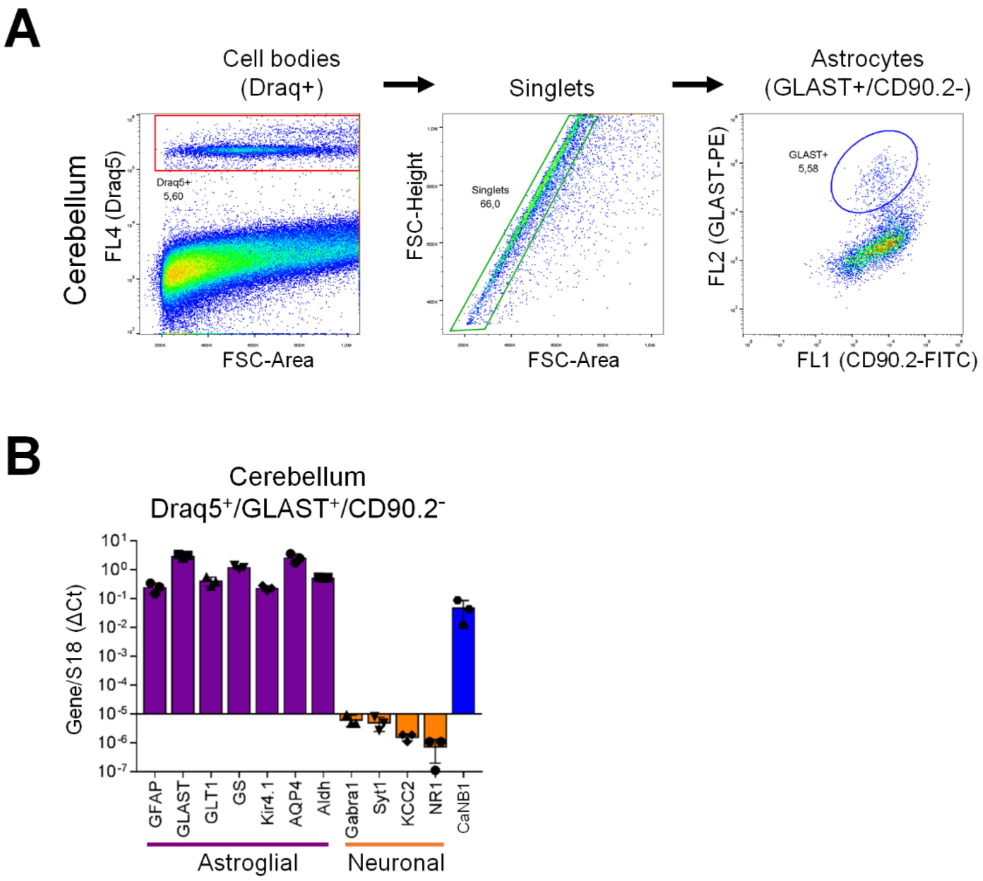


**Supplementary Figure 3. Real-time validation of astroglial CaNB1 KO in FACS separated cerebellar astrocytes. A,** Sorting strategy of astrocytes from P20 ACN-Ctr and KO cerebellum. anti-GLAST (ACSA-1) PE-conjugated antibody was used. **B,** Validation of sorted astrocyte's purity using a panel of primers amplifying genes specific for astrocytes (GFAP, GLAST, Glt1, GS, Kir4.1, AQP4 and Aldh1l1, violet columns) or for neurons (Gabra1, Syt1, KCC2 and NR1, orange columns). Blue columns indicate presence of CaNB1 mRNA in ACN-Ctr astrocytes. In (**A**) a representative experiment is shown. In (**B**) the results are expressed as mean ± SD for 3 independent preparations for each genotype.

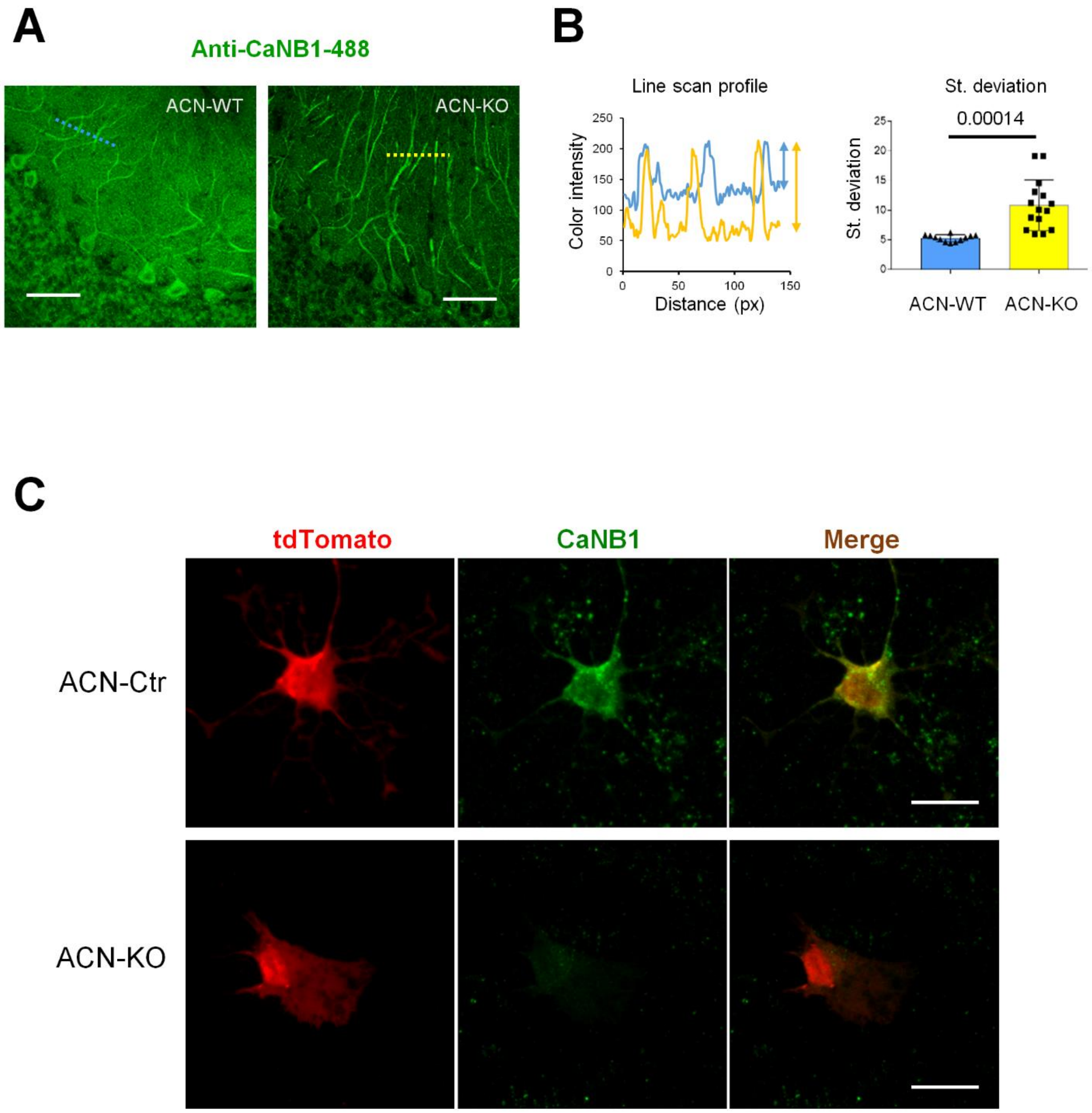


**Supplementary Figure 4. Assessment of CaNB1 KO in ACN-KO mice and RATO-ACN-KO mice. A**, anti-CB1 staining shows high level of CaNB1 expression in cerebellar Purkinje and granule neurons. **B**, line scan profile and significantly increased image standard deviation suggests a non-neuronal reduction of CB1 expression (n = 12 for ACN-Ctr, n = 15 for ACN-KO; mean ± SD). Bar, 90 µm. **C**, confocal images of acutely dissociated hippocampi from P20 RATO-ACN-Ctr (left) and RATO-ACN-KO (right) mouse pups. Note absence of CaNB1 staining (green) in RATO-ACN-KO astrocyte which express tdTomato reporter (red) as compared with RATO-ACN-Ctr astrocyte. Representative images are shown from 3 independent experiments. Bar, 25 µm.

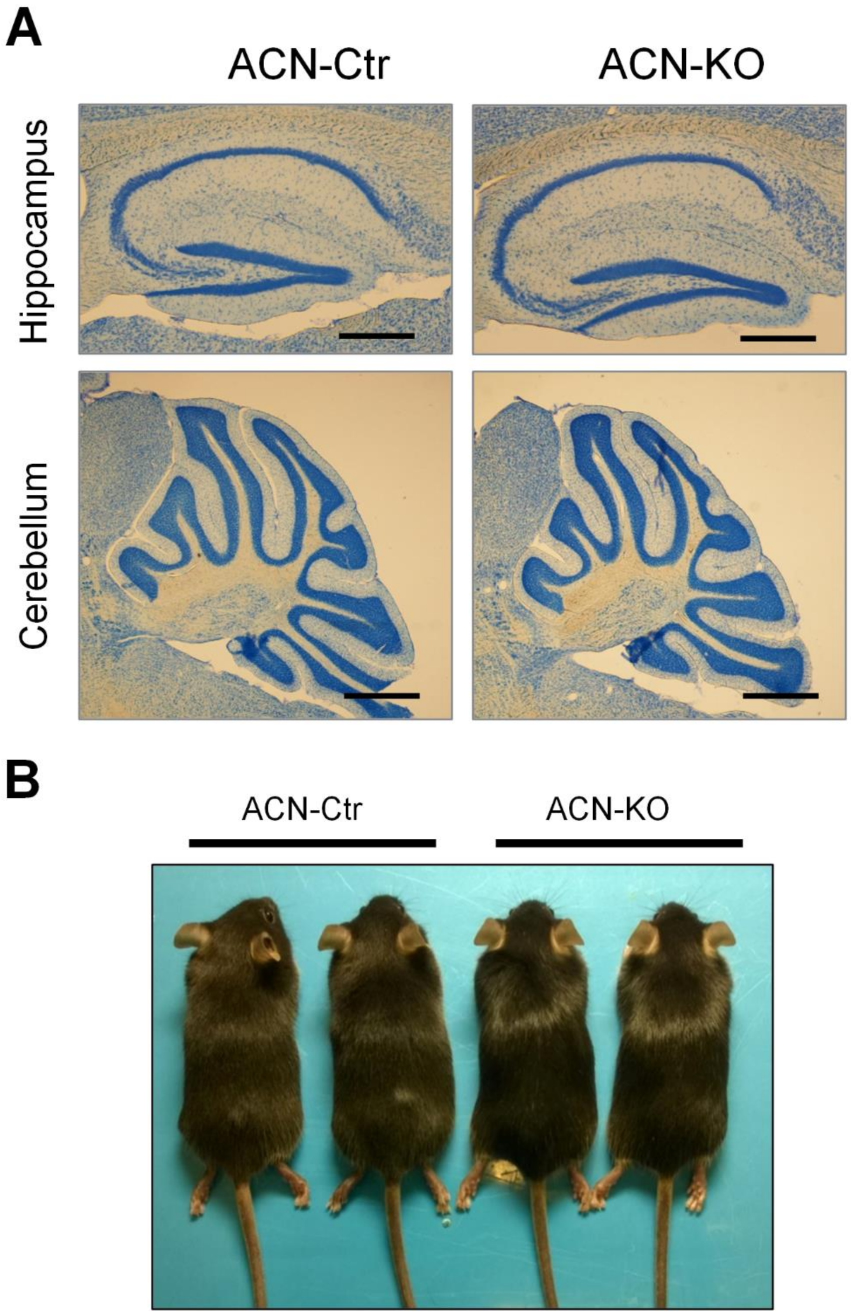


**Supplementary Figure 5. Normal brain development and appearance of ACN-KO mice. A**, Representative Nissle-stained slices of ACN-Ctr (left) and ACN-KO (right) hippocampus and cerebellum. No alterations in cytoarchitecture were found between WT and KO mice. Bar, 100 µM for hippocampus; 500 µM for cerebellum. **B**, Representative photographs of 1 mo old male ACN-Ctr (left) and ACN-KO (right) mice.

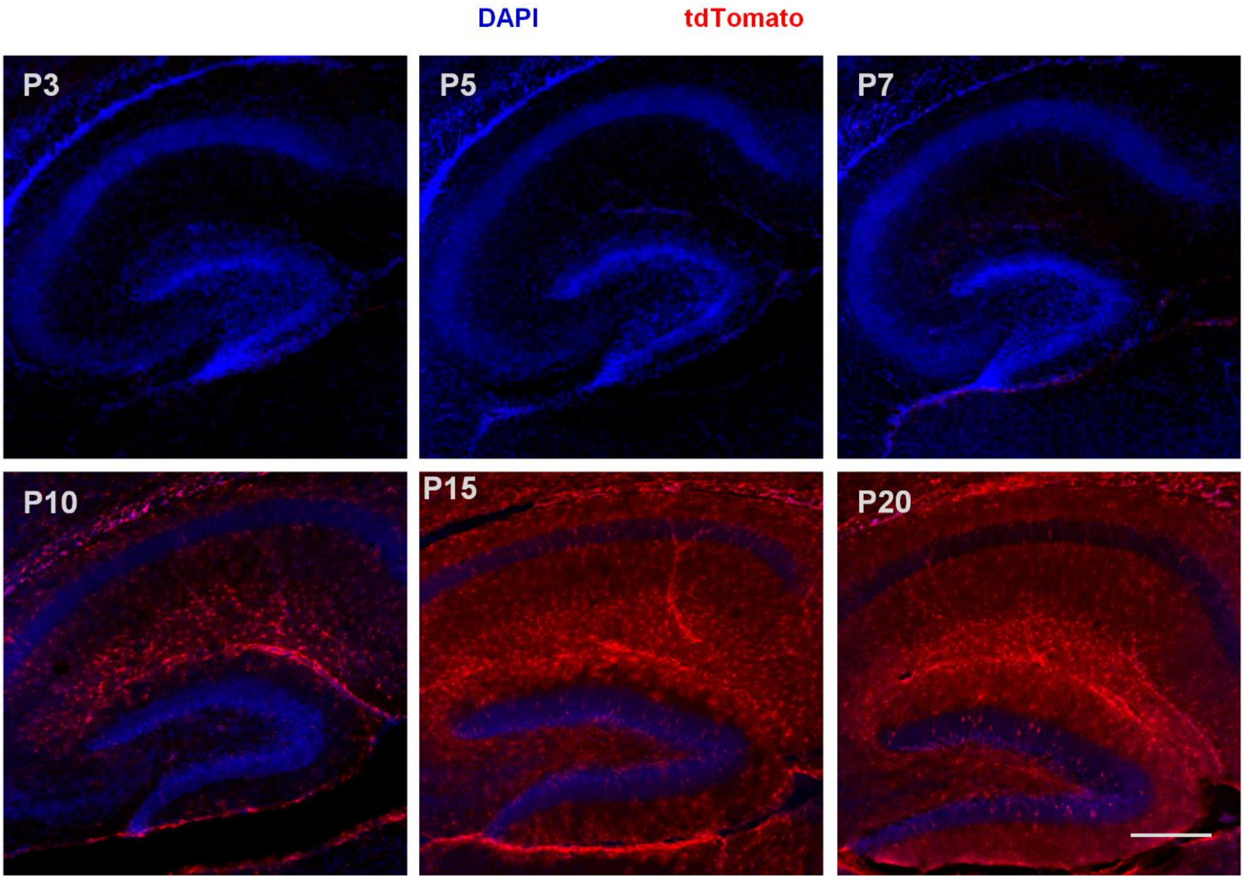


**Supplementary Figure 6. Time course of CaNB1 KO in GFAP-expressing astrocytes in the hippocampus.** Time course of tdTomato expression in the hippocampus of RATO-ACN-KO mice shows appearance of reporter protein between 7 and 10 postnatal days. Full and widespread CaNB1 KO in the hippocampus is achieved by P15-P20. Images are representative from at least 3 ACN-KO mice for each age-point.

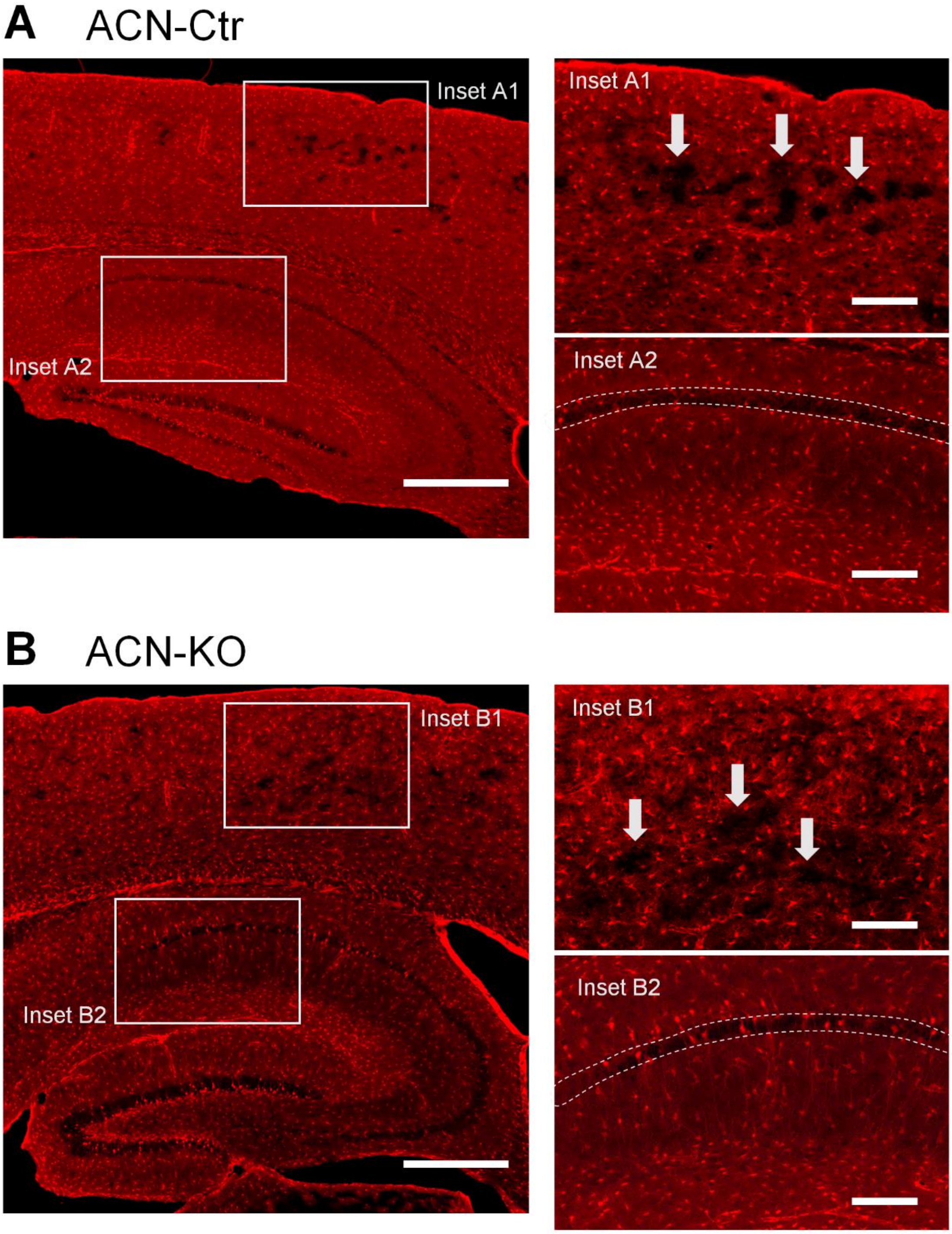


**Supplementary Figure 7. Widespread Cre expression in the hippocampus of ACN-Ctr and ACN-KO mice.** Panoramic confocal z-stack images were taken on sagittal brain sections of ACN-Ctr and ACN-KO mice. Six sections from the brains of three animals per genotype were examined and representative sections are shown. tdTomato fluorescence is shown in red. Note the presence of black areas in the contical regions (arrows) indicating domains of astrocytes not expressing Cre and hence, tdTomato reporter. Note the widespread expression of tdTomato in the hippocampal formation. Bar, 500 μm; bar in insets, 150 μm.

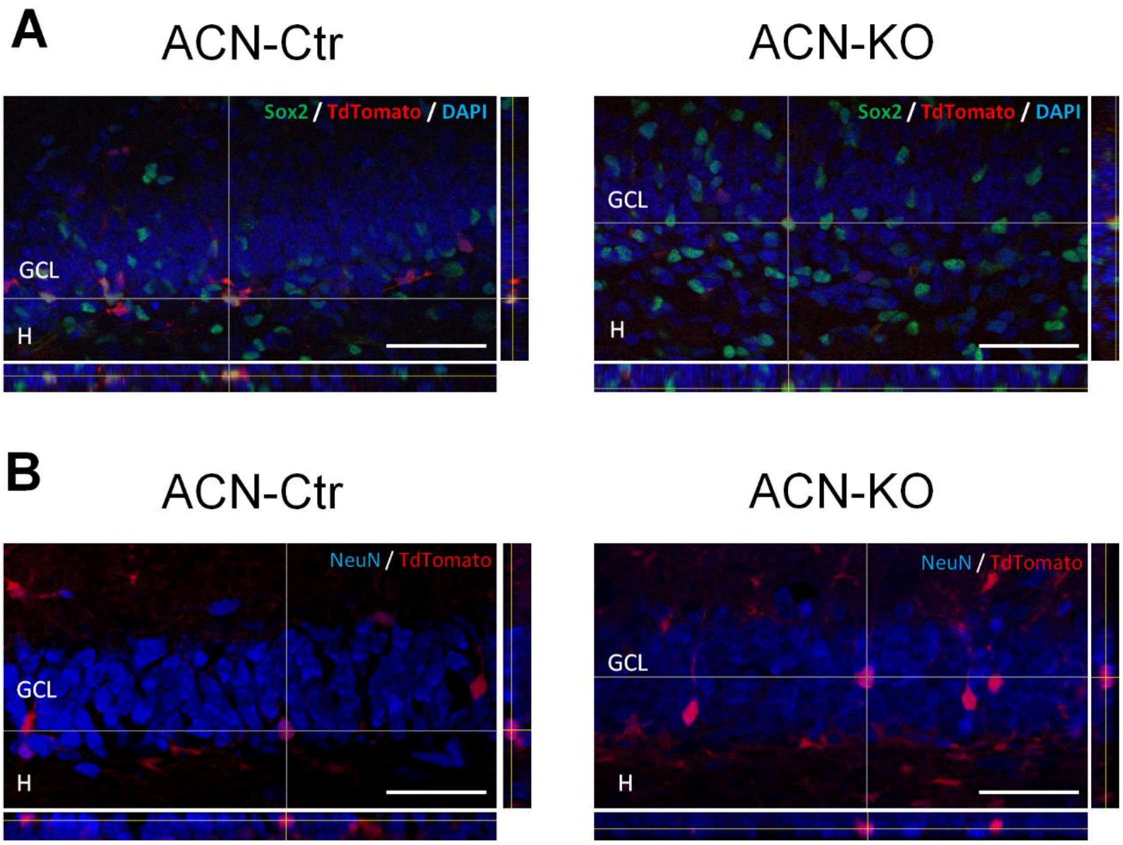


**Supplementary Figure 8. Cre expression does not affect neural progenitors cells and granular mature neurons in the dentate gyrus**. Representative confocal z-plane stack of the dentate gyrus of ACN-Ctr (left pannels) and ACN-KO (right pannels) P7 mice labelled with a Sox2 antibody (**A**) and 1 mo of age mice labelled with NeuN antibody (**B**). Merged images and orthogonal reconstructions showed that at P7, a small amount of Sox2 positive cells (green) also expressed tdTomato reporter (red) in both genotypes (**A**). At 1 mo of age, there were few NeuN-positive neurons (blue) in the granular cell layer expressing tdTomato (red) in both ACN-Ctr and ACN-KO (**B**). These observations may indicate a negligible effect of Cre expression in the neuronal progenitors and in their neuronal progeny. At each time point, one section from the brains of at least two animals per genotype were examined. GCL: granule cell layer; H: hilus. Bar, 50 µm.

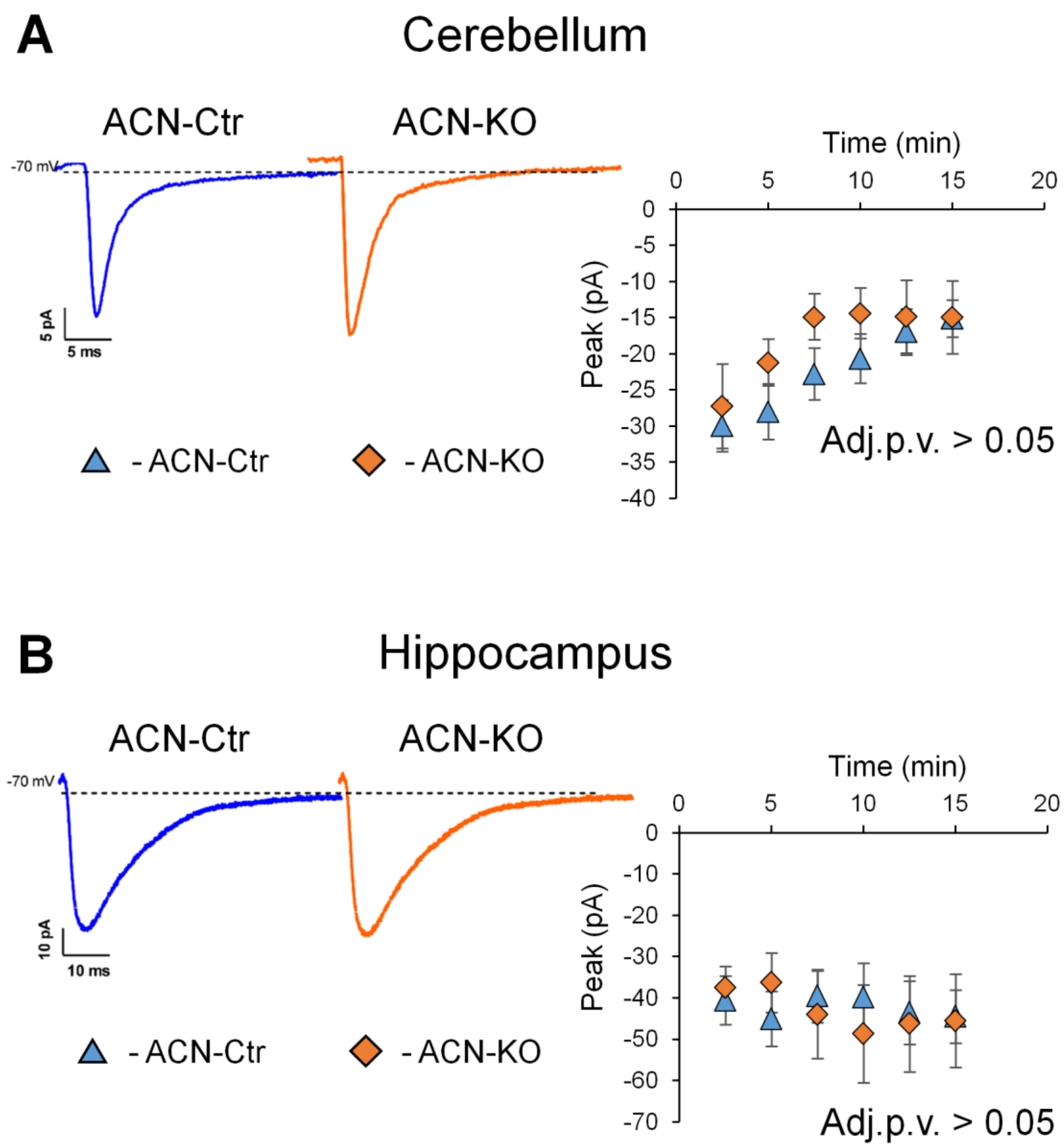


**Supplementary Figure 9. Basal synaptic transmission is not altered in ACN-KO mice. A**, left panel, average of 100 EPSC traces recorded form single cerebellar granule cells in Ctr and KO, following mossy fiber stimulation. Right panel, the histogram shows the average EPSC peak amplitude in Ctr (n = 5) and KO (n = 6). **B**, right panel, average of 100 EPSC traces recorded from single hippocampal CA1 pyramidal neurons in Ctr and KO, following Schaeffer collaterals stimulation. Left panel, the histogram shows the average EPSC peak amplitude in Ctr (n = 5) and KO (n = 6).

CCCCACCTCTTGTTCTCAGGTCTTTCTGGTTTTCCTCCTTTGTTCCTCTCTTGACTCAAATAAGTCCCCCTGTC
TGGATCCCCAGTTAGAGATTAAAGGGCCAGATGTCTACCTCCACTGTGATCTGGGGTCTGAGACAGACTCCCAT
AGGTCCGTGAAGGGAGTGGTCAGGAAGGCGTGGGTGAGAAGCAGGCTGTAGGTGCTCTGCCTGCTGTCACTTGT
TGCCTCTCGAGTACACAGCAGAGCACAGTGTCCTTGGGGCTCTGGTCACTCAAATGCAGGACATGGCAGGGAGC
CCTAGCAGGCTTTGCAAGCAGAGGAAGGGCACAGCCCTTTCCGTTACTGAGACAGCAGAACCAGGCTGTGGGAA
GGCCTGGGCAGTACAAGAAGGTGCCAGCAGGGGTGGGAGGAGGTTGGGGGAGCACTCAGAGCCTTTCTCTTCAG
CCTTACCTTATATCTTGAAGAGGCTAGGGCGATTACTCAGGAACTCTGGCTCAGGCCTATAGATGCTACTTATA
CCCCCCAAAAGGATCATCTATTCACCCCTAAACCCCACACTCCACTAGCCTTCAGGTAAATCTTCTAGAGCCTT
CGTCCCCAGCTTTACATCTTCAGCTGTTCTCTGTGTCCTTGACACACAGGGACTGTGTTTAATTTCCTCTCATG
**ATTTTCC**CAGTCTCATCTACTCCAGACCCAGACCATGAGAATGGACCACAGACCATGAGAAAGTCCATGTGTGC
TTATAGGGCCCATCCTTCCTGCGTAGGCCCAGGAGGAAACAGTTAGGGGCAGGTAGGCGCTGACCCTCCGGTGT
CTGTAAT**GGAAAAAT**CTCAGCTGCAGAGCCGGAGGTGGCCAGCTATTTAGGGAATTCAGATGTGAGCCCTGTGC
TCTGAGAGTAGATTTCCTGGACAAGAGGTCCATAATCGCTCTACAGGGAGCTCCCTCCAGGGCCAGAGGGGGCT
CAACCCAGCCAGCCCAGCCAGCGGGCACCTTAGGACTCCTGCCAATGCCTCATTGCCTCCCTCCTCCTTCCCAG
CCAGGGTTCTCAGACCTGCCCCTGCTCAGTGGGCGTCCACTCACTCACTACATTTCGGTGCCCAGAGAAATGGG
GACACTGCAGAACTTTATCCCTCTAACTGTTCCTAGTCGGTGGGGGCAGAAAATCTTCTTCTCTCTGTCCCCAG
CCTCCATTCTTCAACAAATGAATGATGCCAACACCCCCCCCGCCCCCAGAGCGGCTGGGCTGGTCATGAGTTA
TCTGCTCTGTTCCTGTTTCTATTGAGAGTAATGTTGAAGACATCTTCTCAGCTGGAGGTGAAGCCCCCTCCCTC
CCCCCAGACCCAACCTCCAGGATCTCCCATGCCATTAAACAGGTCTCTCACAGTGGAGTCTGCCTGACCCCTTC
CTCCCAGAAGGAACCAAGGAAGTCCTTGCTCTTTCTAGTCCCACAGAGTTGGTATTTGAGGTGATAATGACTTC
CATGCCCTCCTGCTACACCTTCCAGATTTGACTTCTCATACCATTCACCGACCTCCGACTCAGACTCACTCAAA
GCTCCCCTGTAGCAGTGAAGGGGAGGCTCATCTGCTTCATACACAGAATGGCAGAAATGCTTCCACGTGCCTGT
TGTGCCCGGACTTGGGCGGCCTTCCTGGCTCTTTCATATCTTCTGTCCTTCACCTTCCCAACTGCTTCTAAGTG
CGGAGACTCTTGCTTGTCTTGAACACTGGGTTACAGGTGTGTCCACCATTAAAAATCATGTCACAATGTTATAA
CCAACAACATTCATGTCATCTACAAACCACACTGGACAACTATGAAATCACATTATCCAGAGAGAGAGAGAAAG
AGAGAGAGAGAGAGAGAGAGAGAGAGAGAGAGAGAGAGAGAGAGAGAGAGAGAGAGAGAGAACGAACGTTGGTGGT
GGCACATTCTCTTAATCCCAGCAGAGGCAGGGGGATCTCTGAATTTAAGGATTAGCCAGATCTGTAGAGCGATT
TCCACAACAGCCAAAGCAACACAGAGAAAACTTGTGTCAAAAAACAAACAAACAAATAAAAAATCGCTACACTG
TTACAAAGCATCTCAATGTTTGAAGTAAGGTTAGGATGTATTGGGCCAAATTCCTAGCTATGGTGGTGGTGGGG
TGGCGTATGGAGCCCAAGAGCAACAAGTGGGACATGCCTGGTGCAGCATCTCCAGGCTTTGGTTCTGCAAGGAG
CTTCAAAAGCAAGAGTCGTCTGATTCTAGTTCTAATTCTTTCTTGGTCACAACGTCCTTGCAGCCCCTTACAGC
AGATGGAGCAGGCTGTTGCTTTCTCTGGAAACTTGGATGTGAGCTCAGGGAGATTCTGTCTTATCCAGTAAGAT
TGCTGAATTCTTATAGCCTTGCTTCCCCACCT**GGAAATAT**AGGCAATTAGTCCCAGGACCATGAGCTGGCTGCT
CTCGAGGGGCCACCTCTCC**TTTTTCC**AGCTACCCCTTCCCCATTCCTTTCCCCTTGCTTCTCTCTTGGTTGTAG
CTGCCCATGGGATGGGGAAATGCTAAGAGGAGCTGAGGGGGACAGACCGGGGACAAAACAGGACCATCAGCTGG
GCCAGTTGCTAAAGAGGCGGGAGGGGAGGGGAGGAGTCCTCAGGGATCCAGTTTCAACAAACGTTTCTTTCCCC

AGAAGGGGAAGGCGGGAGTGAGGGGGAGAGGGACCTATTTAAAGCTACCCTGTTGCTCAGACTGTCTCTGTCTG
TCTGCCAGGGTCTCCAGCTGCCCCAGACAGGCGGTGTGGTCTTGGGATCCTCCTGGTGACCTTTCCAGCCTAGG
TCCCCTCAGCCACTCTGCCCCAAG**ATG**GGTCGTGGGGTGAG

**Supplementary Figure 10. In silico analysis of NFAT-binding sites in promoter region of Atp1a2 gene.** NFAT-binding sites were found in distal (green, -1843 bp and -1998 bp) and in proximal (red, -203 bp and -264 bp) regions of Atp1a2 promoter region. Transcription start site is indicated by an angled arrow and mRNA region is colored in blue.

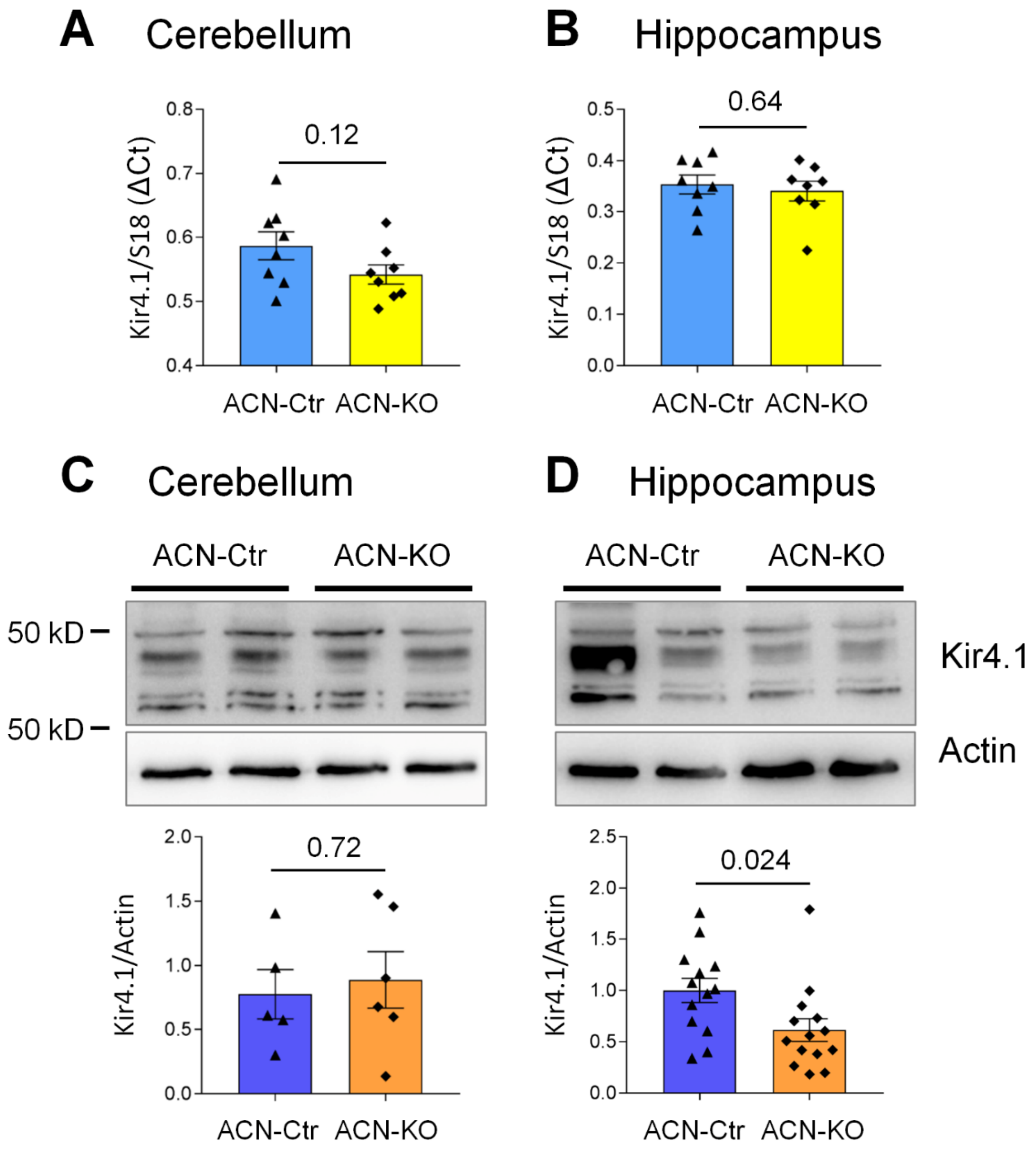


**Supplementary Figure 11. Kir4.1 protein is downregulated in the ACN-KO hippocampus.** Real-time PCR relative mRNA quantification of Kir4.1 in whole cerebellar (**A**) and hippocampal (**B**) tissues of ACN-Ctr and ACN-KO mice. Data are expressed as mean ± SEM, n = 8 mice per genotype. Western blot analysis of protein expression of $K_{ir4.1}$ in cerebella (**C**) and hippocampi (**D**) of ACN-Ctr and ACN-KO mice. Data are expressed as mean ± SEM, n = 5 mice for ACN-Ctr, n = 6 mice for ACN-KO in the cerebellum. In the hippocampus n = 13 for ACN-Ctr, n = 14 for ACN-KO.

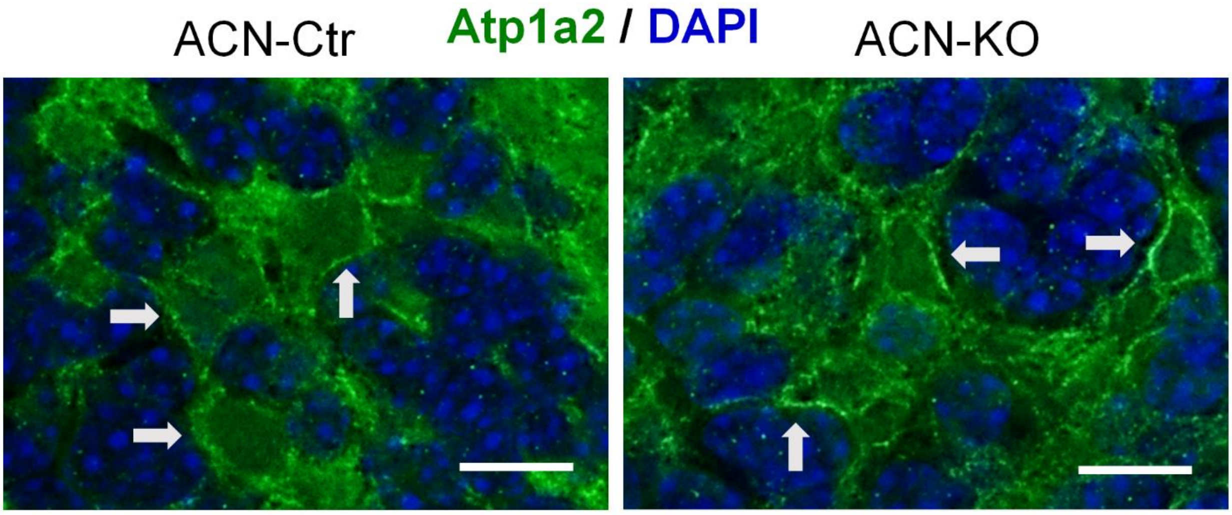


**Supplementary Figure 12. Expression and localization of Atp1a2 NKA subunit is not altered in the cerebellum of ACN-KO mice.** Expression of Atp1a2 subunit of NKA was detected in granular layer glial cells of the cerebellum of ACN-Ctr (left panels) and ACN-KO (right panels) mice (green). Nuclei were counterstain by DAPI (blue). Images are representative for 6 sections per brain from three mice per genotype. Bar, 15 µm.

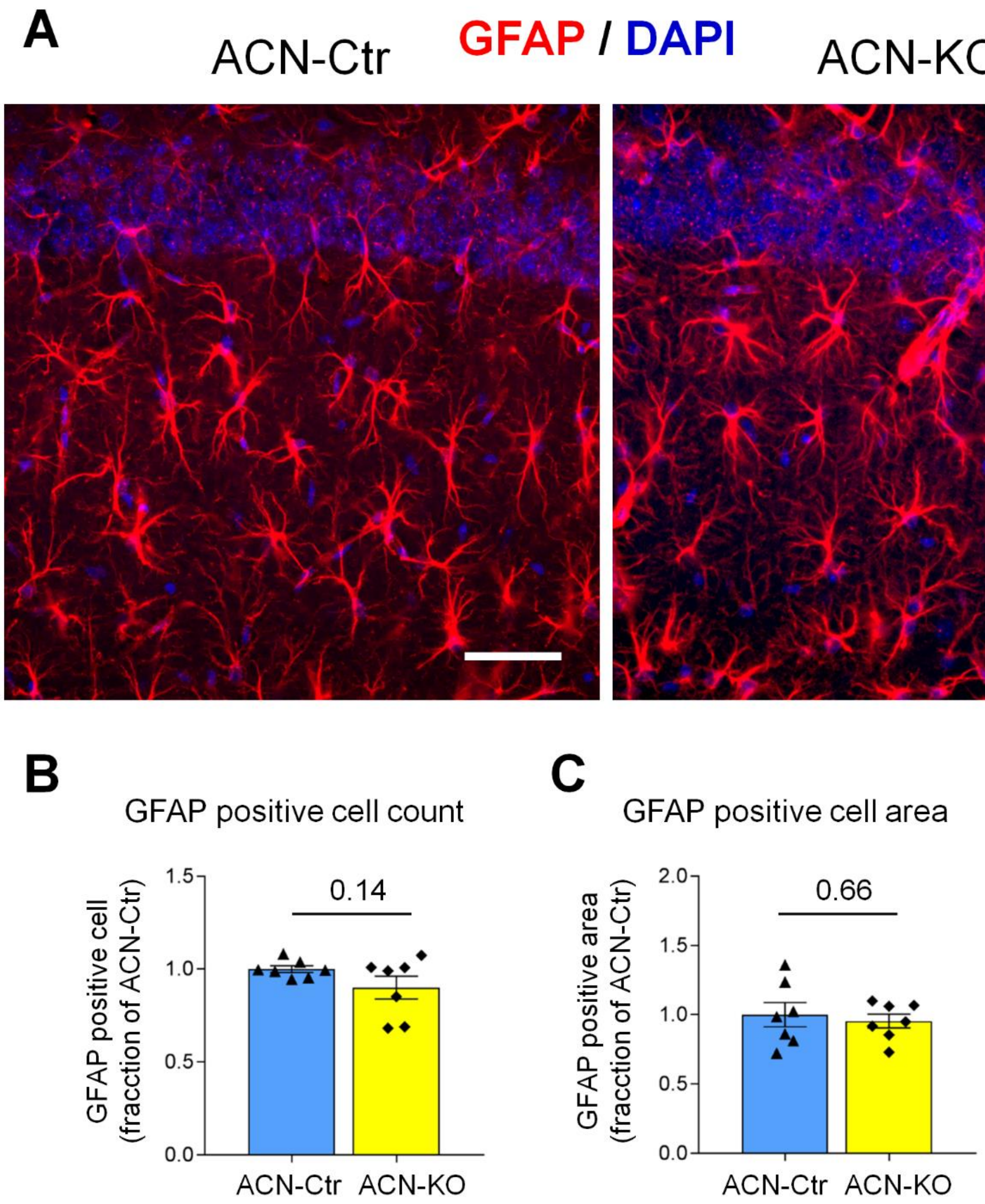


**Supplementary Figure 13. Number and area of GFAP positive cells is not different in the hippocampus of ACN-Ctr and ACN.KO mice. A**, immunofluorescence analysis of GFAP-positive astrocytes in hippocampus of ACN-Ctr and ACN-KO mice. A fragment is shown corresponding to CA1 region. Fraction of GFAP-positive cells (**B**) and GFAP-stained area (**C**) are not different between ACN-Ctr and ACN-KO mice. Data are expressed as mean ± SEM of seven animals per genotype. Bar, 50 µm.

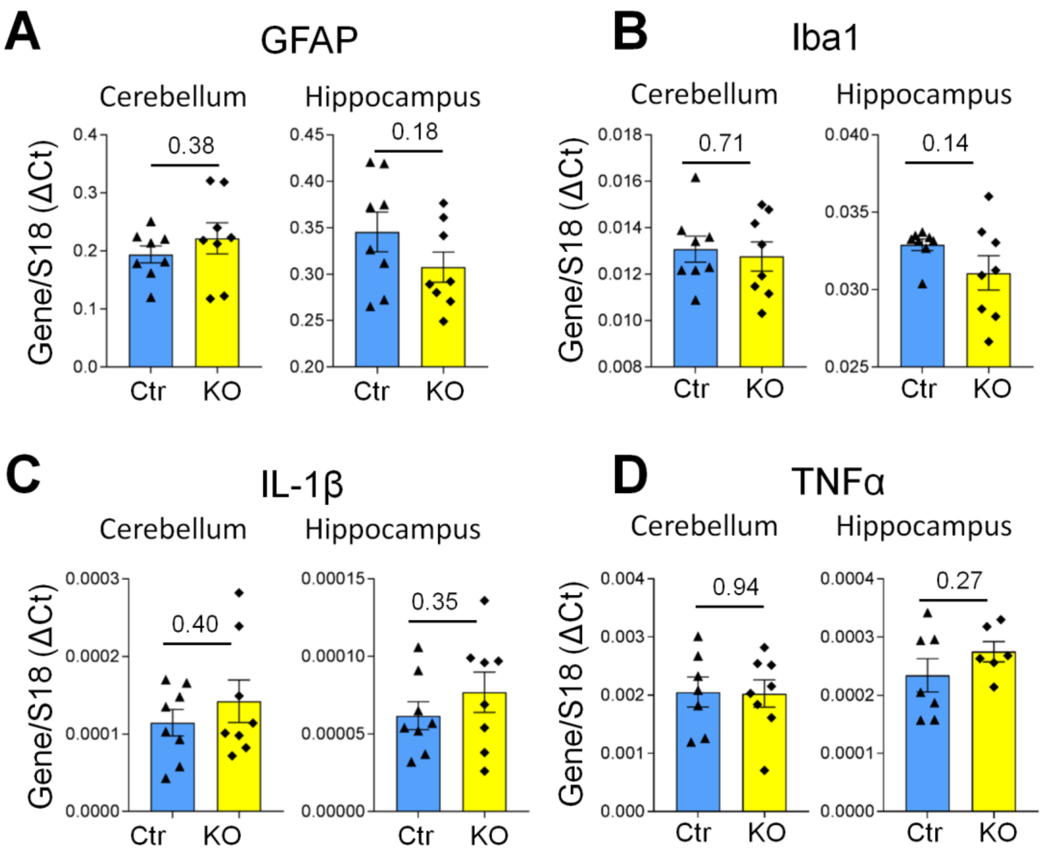


**Supplementary Figure 14. Expression of astroglial markers and inflammatory genes is not changed in ACN-KO.** Real-time PCR analysis of mRNA levels of GFAP (**A**), Iba1 (**B**), IL-1β (**C**) and TNFα (**D**) in the cerebellum and the hippocampus of ACN-Ctr and ACN-KO mice. Data are expressed as mean ± SEM of 6-8 mice per genotype.
